# Phylogenetic Dissection Provides Insights into the Incongruity in the Tree of Archaeplastida Between the Analyses of Nucleus- and Plastid-Encoded Proteins

**DOI:** 10.1101/2025.05.06.652364

**Authors:** Ryu Isogai, Ryo Harada, Takuro Nakayama, Yuji Inagaki

## Abstract

Archaeplastida is defined as a taxonomic assemblage comprising three sub-clades, namely Chloroplastida, Glaucophyta, and Rhodophyta plus two non-photosynthetic lineages sister to Rhodophyta (the latter three lineages collectively termed “Rhodozoa” here). The members of Archaeplastida are the descendants of the eukaryote that took up and transformed a cyanobacterial endosymbiont into a primary plastid. Recent phylogenetic analyses of multiple proteins (phylogenomic analyses) stably recovered the monophyly of Archaeplastida, but uncertainty remains in the relationship among the three sub-clades in this assemblage. The phylogenomic analyses of nucleus-encoded proteins (nuc-proteins) grouped Chloroplastida and Glaucophyta together, excluding Rhodozoa in the Archaeplastida clade, albeit the union of Chloroplastida and Rhodophyta was often inferred from the phylogenomic analyses of plastid-encoded proteins (pld-proteins). In this study, we challenged the previously recognized but as-yet-explicitly addressed issue in the tree of Archaeplastida (ToA). The detailed analyses of the nuc-protein and pld-protein supermatrices revealed that taxon sampling can invoke different types of phylogenetic artifacts into the inferences from both supermatrices examined here. In the end, we propose a working hypothesis for the ToA and provide future perspectives toward resolving the ToA.

## Introduction

The plastids surrounded by double membranes have been found in three photosynthetic lineages, namely Chloroplastida, Glaucophyta, and Rhodophyta (Adl et al. 2005). This type of plastid, so-called “primary plastids,” was most likely established through “primary endosymbiosis” between a cyanobacterium (endosymbiont) and a heterotrophic eukaryote (host) (Archibald 2009, Keeling 2013, Sibbald and Archibald 2020). It is generally believed that transforming a cyanobacterial endosymbiont into a plastid governed by the host eukaryotic cell is rare in the Tree of Life (ToL). Thus, Chloroplastida, Glaucophyta, and Rhodophyta have been anticipated to form a monophyletic assemblage (i.e., a supergroup Archaeplastida), evolving from a common ancestor, the first eukaryotic phototroph bearing a primary plastid (Archibald 2009, Keeling 2013, Sibbald & Archibald 2020). Nevertheless, recent studies updated our understanding of the organismal diversity of Archaeplastida, as well as the rarity of primary endosymbiosis in the ToL. First, two non-photosynthetic groups, Rhodelphidia and Picozoa, were reported to belong to Archaeplastida (Gawryluk et al. 2019, Schön *et al*. 2021). Phylogenetic analyses based on multiple proteins (phylogenomic analyses) placed both of them at the base of Rhodophyta (note that we tentatively name the clade of Rhodophyta, Rhodelphidia, and Picozoa as “Rhodozoa” in this study). Primary plastids can be considered as a convincing plesiomorphy uniting Chloroplastida, Glaucophyta, and Rhodophyta (Rodríguez-Ezpeleta et al. 2005) (representative algal species are shown in Fig. 1a). The data from Rhodelphidia suggested that they still retain the remnant plastid (Gawryluk et al. 2019), not violating the ancestral acquisition of a primary plastid in Archaeplastida. On the other hand, no strong evidence for secondary loss of an ancestral plastid has been found in the picozoan genome data, implying either the complete loss or the primary absence of a plastid in this lineage (Schön *et al*. 2021). If the latter is the case, the plastids in rhodophytes and rhodelphids and those in chloroplastids and glaucophytes could have been evolutionarily distinct from one another. So far, no plastid-encoded protein (pld-protein) is available from any picozoan or rhodelphid, we do not address the uncertainty in Picozoa described above and assume the monophyly of the plastids in Archaeplastida in this study. Apart from Archaeplatida, we now know the second case of the acquisition of a photosynthetic organelle derived from a cyanobacterium. Testate amoeba belonging to Rhizaria (*Paulinella* spp.) appeared to bear photosynthetic organelles that are evolutionarily distinct from but equivalent to primary plastids in Archaeplastida (Bhattacharya et al. 1995; Keeling 2004; Marin et al. 2005; Yoon et al. 2006; Nowack et al. 2012; Nowack et al. 2016; Singer et al. 2017; Macorano and Nowack 2021). The host and endosymbiont phylogenies of the eukaryotes bearing cyanobacterium-derived photosynthetic organelles with double membranes claim that primary endosymbiosis occurred (at least) two separate branches of the ToL (Stephens et al. 2021).

**Figure 1.**
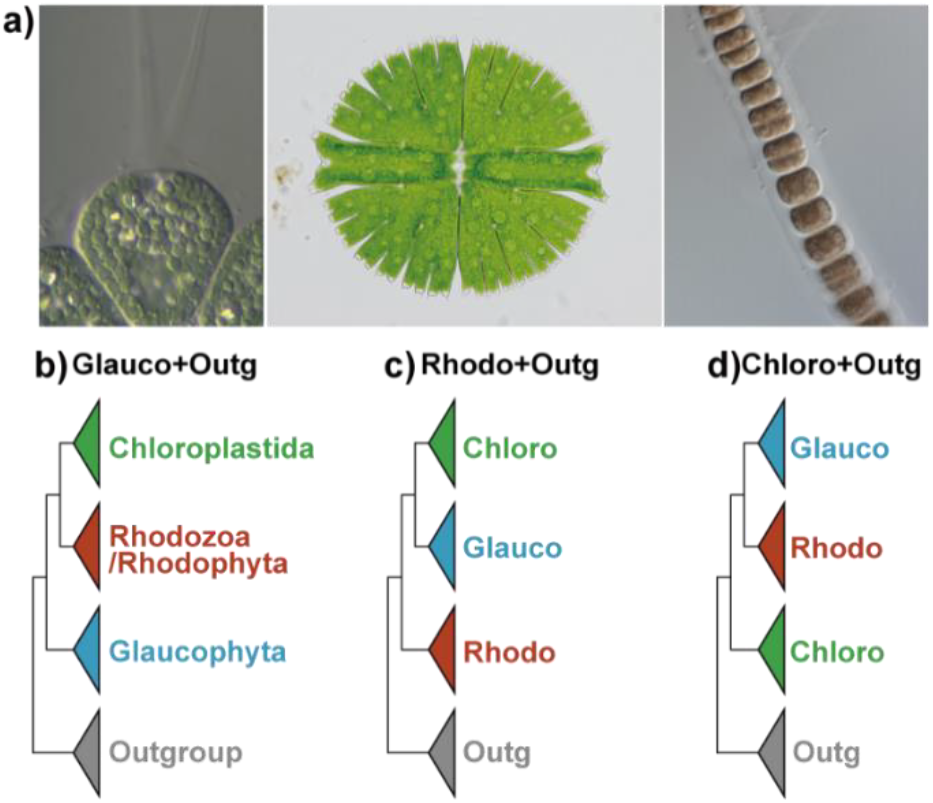
Algal species representing the subclades of Archaeplastida and three types of ToA. (a) The glaucophyte *Gloeochaete* sp. (left), the green alga *Micrasterias* sp. (center), and the red alga *Stylonema* sp. (right). The images were provided by Dr. Takeshi Nakayama (University of Tsukuba, Japan). (b) “Glauco+Outg” tree, in which Chloroplastida and Rhodozoa/Rhodophyta are connected, resulting in the sister relationship between Glaucophyta and the outgroup. (c) “Rhodo+Outg” tree, in which Chloroplastida and Glaucophyta are connected, resulting in the sister relationship between Rhodozoa/Rhodophyta and the outgroup. (d) “Chloro+Outg” tree, in which Rhodozoa/Rhodophyta and Glaucophyta are connected, resulting in the sister relationship between Chloroplastida and the outgroup. Chlorophyta, Glaucophyta, and Rhodozoa/Rhodophyta are abbreviated as Chloro, Glauco, and Rhodo, respectively, in **c** and **d**.

Prior to incorporating the data from recently found, non-photosynthetic relatives of Rhodophyta, as well as a novel species related to Cryptista, into phylogenomic analyses, the host monophyly of Chloroplastida, Glaucophyta, and Rhodophyta had been challenging to recover in phylogenomic analyses until recently (Mackiewicz and Gagat 2014). In phylogenomic analyses of nuc-proteins, the monophyly of Archaeplastida was often hindered by Cryptista comprising cryptophyte algae and their non-photosynthetic relatives (goniomonads, kathablepharids, and Palpitomonas bilix) (Burki et al. 2016, Gawryluk et al. 2019, Strassert et al. 2019). Fortunately, Archaeplastida was reconstructed as a clade with confidence after carefully assessing the proteins and taxa to be included in phylogenomic analyses (Yazaki et al. 2022, Irisarri et al. 2022). Irisarri et al. (2022) examined which proteins were to be (or not to be) included in a supermatrix to recover the monophyly of Archaeplastida. Yazaki et al. (2022) demonstrated that the taxa branched at the base of Rhodophyta (e.g., Rhodelphis spp.), and those positioned at the base of and sister to Cryptista (e.g., Palpitomonas bilix and Microheliella maris) were critical to suppress the putative false phylogenetic affinity between Rhodophyta and Cryptista and led to the recovery of the monophyly of Archaeplastida. The phylogenomic analyses, in which the suspicious affinity between Rhodophyta and Cryptophyta was suppressed efficiently, further nominated the assemblage of M. maris and cryptists (collectively designated Pancryptista) as the closest lineage of Archaeplastida (Yazaki et al. 2022).

We are now comfortable about the monophyly of Archaeplastida, but there is still an unsettled issue in the tree of Archaeplastida (ToA)—the phylogenetic relationship among Chloroplastida, Glaucophyta, and Rhodozoa remains controversial. In theory, three types of ToA, depending on which two out of the three sub-clades in Archaeplastida form a sister relationship, are possible (Figs. 1b-d). Traditionally, Glaucophyta was hypothesized to emerge from the last common ancestor of Archaeplastida prior to the separation of Chloroplastida and Rhodophyta, corresponding to the “Glauco+Outg” tree shown in Fig. 1b. This is mainly due to a ‘cyanobacterium-like’ characteristic that was thought unique to the plastids of glaucophytes (glauco plastids) until recently. The glauco plastids are surrounded by the peptidoglycan wall and were regarded as an evolutionary intermediate bridging plastids and free-living cyanobacteria—the former usually lack and the latter bear peptidoglycan walls (Löffelhardt and Bohnert 2001). More recent studies found plastids with peptidoglycan walls in restricted members belonging to Streptophyta, a subgroup of Chloroplastida, e.g., the charophyte alga Klebsormidium nitens (Takano et al. 2018), the moss Physcomitrella patens (Hirano et al. 2016), and the lycophyte Selaginella moellendorffii (Takano and Takechi 2010). Accordingly, the updated distribution of peptidoglycan-surrounded plastids demands multiple losses of the particular trait in Archaeplastida and lends no strong support for any type of ToA. Likewise, no conclusive picture of the ToA has been drawn from molecular-based analyses conducted to date. Previously published nuc-protein-based phylogenomics often grouped Chloroplastida and Glaucophyta, recovering the sister relationship between Rhodozoa/Rhodophyta and other eukaryotes (outgroup) with high statistical support (Tice et al. 2021; Tikhonenkov et al. 2022; Eglit et al. 2024). Henceforth, this type of ToA is designated as “Rhodo+Outg” tree (Fig. 1c). We can also examine the internal branching patterns in the ToA by analyzing pld-proteins if the plastids in the extant members of Archaeplastida have been inherited vertically from their common ancestor. To our knowledge, there has been little ground to doubt the vertical inheritance of plastids in the extant photosynthetic members in Archaeplastida, and thus the ToA inferred from pld-proteins is supposed to agree with, or at least does not conflict fundamentally with, that from nuc-proteins. In reality, the ToA with the union of Rhodophyta and Chloroplastida—Glaucophyta and the outgroup (cyanobacteria) tied directly (Glauco+Outg tree; Fig. 1b)—was typically inferred from multiple pld-proteins (Rodríguez-Ezpeleta et al. 2005, Figueroa-Martinez et al. 2019). In the third type of the ToA, the “Chloro+Outg” tree, Rhodozoa/Rhodophyta and Glaucophyta group together, excluding Chloroplastida (Fig. 1d; this tree has been rarely recovered in past phylogenomic studies). Unfortunately, the uncertainty in the phylogenetic relationship among the three subclades of Archaeplastida has not been explicitly addressed in any past study.

There are two general approaches to improving organismal phylogeny. The first approach is a computational approach that remediates the methodological issues in tree reconstruction. The second approach is more biological than the first and searches for novel lineages/species in nature, incorporating them into phylogenetic inferences. Although the two approaches are not mutually exclusive, we took the computational approach to address the issue in the ToA in this study, as novel species belonging to Archaeplastida have not been reported after Rhodelphidia and Picozoa. We here conducted phylogenomic analyses of an alignment comprising 276 nuc-proteins (nuc276) and that comprising 54 pld-proteins (pld54) to understand the reasons why the ToA inferred from the two types of multi-protein alignments (supermatrices) conflicted with each other. Our detailed assessments revealed that taxon sampling severely affected the phylogenetic inferences from both nuc276 and pld54. Depending on taxon sampling, fast-evolving positions and heterotachious positions appeared to overrate a particular type of the ToA in the nuc276 and pld54 analyses, respectively. Finally, we propose a type of ToA as a working hypothesis and reinforce the potential significance of as-yet-discovered members of Archaeplastida, particularly those branched at the base of the three sub-clades, to infer the genuine ToA in future studies.

## Materials & Methods

### Nuclear Protein-Based Phylogenomic Analyses

#### Supermatrices

We modified 351 single-protein alignments considered in a previous phylogenomic analysis (Harada et al. 2024). The single-protein alignments were updated by adding the sequences of 25 species belonging to Archaeplastida and three operational taxonomic units (OTUs) for Picozoa (see below in this section; Schön et al. 2021). Overall, the above-mentioned archaeplastid species represent most of the classes and/or orders in Archaeplastida proposed in Adl et al. (2019). Separately, we updated the sequence data of the green alga Mesostigma viride in this study, as the coverage of this species was poor at approximately 17.56% in the original supermatrix. For 20 archaeplastid species, the amino acid sequence data were retrieved from the OneKP database (Leebens-Mack et al. 2019; Carpenter et al. 2019). For six archaeplastid species (including M. viride) that are not available in the OneKP database, their transcriptome data were obtained from the NCBI Sequence Read Archive (SRA). The nucleotide sequence data were assembled into contigs using Trinity ver. 2.13.2 (Grabherr et al. 2011) after quality control by fastp ver. 0.23.2 (Chen 2023) with options ‘-q 20 -u 80.’ The contig (nucleotide) sequences were translated into amino acid sequences using TransDecoder.LongOrfs ver. 5.5.0 (https://github.com/TransDecoder/TransDecoder) with an option ‘-m 50.’ Schön et al. (2021) generated the single amplified genomes (SAGs) and COSAGs from the picozoan cells and created three OTUs. We followed the procedure taken by Schön et al. (2021) to generate the three picozoan OTUs by merging (i) the amino acids data of SAG22, SAG31, and SAG11, (ii) those of COSAG02, COSAG03, COSAG04, and COSAG06, and (iii) those of COSAG05, COSAG01, and SAG25. The amino acid sequence data considered additionally or updated for assessing the ToA (29 archaeplastid species/OTUs in total; see described above) were subjected to CD-HIT ver. 4.8.1 (Fu et al. 2012; Li and Godzik 2006) with an option ‘-c 0.98’ to reduce the sequence redundancy and subsequently formatted into the databases for blastp (see below).

Besides improving taxon sampling in Archaeplastida (see above), we added the amino acid sequence data of the two Anaeramoeba species, which were provided as supplementary data in Stairs et al. (2021), to update the general diversity of eukaryotes in the supermatrix prepared in this study. The Anaeramoeba sequences were also processed by CD-HIT as described above and then formatted for blastp.

The 351 nuc-proteins, which comprised the supermatrix analyzed in Harada et al. (2024), were searched for individually in the 31 blastp databases (see above). We ran blastp ver. 2.12.0+ (Camacho et al. 2009) against each database with all the sequences included in the previous single-protein alignment as queries. The blastp options were set as ‘-evalue 1e-30 - max_target_seqs 5 -max_hsps 1.’ The candidate sequences retrieved by blastp were aligned with the corresponding alignment by MAFFT v.7.490 (Katoh and Standley 2013), followed by trimming the positions occupied by a large number of gaps and those aligned ambiguously by trimAL v. 1.4.rev15 (Capella-Gutiérrez et al. 2009). Then, the updated alignments were subjected to the maximum likelihood (ML) phylogenetic analyses by using IQ-TREE v. 2.1.2 (Minh et al. 2020). We individually inspected the resultant ML trees to identify and discard (i) potential paralogs, (ii) potential sequences horizontally acquired from organisms distantly related to Archaeplastida, and (iii) “stand-alone” long branch sequences. (i) We regarded the ‘archaeplastid sequences,’ which formed a clade separated by a long branch from other sequences included in the original alignment, as potential paralogs. (ii) If an ‘archaeplastid sequence’ and that of a eukaryote (or those of eukaryotes) distantly related to Archaeplastida were grouped with high statistical support, such sequence was considered to be horizontally transferred. (iii) In addition, we occasionally observed a long branch ‘archaeplastid sequence,’ which showed no affinity to any other sequences analyzed together, in the ML tree. Such “stand-alone” long branch sequences can be (a) paralogs that were included solely in the particular alignment, (b) horizontally acquired, albeit their potential donors were absent in the alignment, or (c) bacterial sequences contaminated. The above-mentioned three types of ‘archaeplastid sequences’ can bias phylogenetic inferences of the organismal relationship among Chloroplastida, Glaucophyta, and Rhodozoa/Rhodophyta. Thus, we omitted them from the downstream analyses described below. The remaining sequences after the operations described above were realigned, and then the alignment positions were reselected for the phylogenetic analyses described below. Note that we observed amino acid sequence stretches, which were highly diverged compared to other orthologous sequences, in some picozoan proteins. These regions likely emerged by erroneous translation of introns into amino acid sequences, as Schön et al. (2021) applied an annotation program designed for prokaryotic (intron-lacking) genomes to the picozoan genome data. Such regions are problematic for phylogeny but can remain in alignments after the trimming procedure applied in this study. Thus, the final supermatrix contains two single-protein alignments in which we replaced the amino acid sequence stretches likely derived from translating intron (nucleotide) sequences in picozoan proteins with “X,” a character representing an uncertain amino acid identity (134_ODO2A_full_mafft.fasta and 260_nsf1-I_full_mafft.fasta; see the supplementary data).

Before the concatenation of the selected single-protein alignments into the final supermatrix, we conducted the third series of aligning and trimming of the sequences by MAFFT with the L-INS-i algorithm and the combination of trimAL with ‘-gt 0.1’ option and BMGE ver. 2.0 (Criscuolo and Gribaldo 2010) with ‘-g 0.3 -e 0:0.6 -w 5’ options. We concatenated the 345 single-protein alignments into a preliminary supermatrix. After omitting (i) the OTUs, in which gaps occupied more than 80% of alignment positions, and (ii) proteins that were absent in more than 40% of the OTUs from the initial supermatrix, we obtained a supermatrix comprising 317 proteins from 133 OTUs (nuc317; 117,125 amino acid positions in total). nuc317 was used to infer the global eukaryotic phylogeny. Next, as we were primarily interested in the ToA, a smaller supermatrix, “nuc276,” in which 276 single-proteins (94,907 amino acid positions in total) were sampled from 57 OTUs, was generated from nuc317. nuc276 comprised 25 species of Chloroplastida, four species of Glaucophyta, 16 species of Rhodozoa (11 species of rhodophytes, two rhodelphids, and three picozoan OTUs), and 12 species of other eukaryotes as the outgroup. The outgroup in nuc276 included six species in Pancryptista, which was nominated as the closest related lineage of Archaeplastida (Yazaki et al. 2022), but any photosynthetic member (e.g., cryptophyte algae) was omitted to avoid biasing phylogenetic inferences due to genes potentially transferred from the photosynthetic endosymbionts that gave rise to the current plastids. The single-protein alignments, of which more than half the number of the species in each of the three subclades of Archaeplastida—(i) Chloroplastida, (ii) Glaucophyta, and (iii) Rhodozoa, were retained in nuc276. In every step preparing nuc276 from the preliminary supermatrix (see above), the single-protein alignments were individually realigned by MAFFT with L-INS-i algorithm and then subjected to a two-step trimming of ambiguously aligned positions and those occupied by gaps, initially by trimAl with ‘-gt 0.1’ option and then by BMGE with ‘-g 0.2 -e 0:0.5 -w 5’ options.

The supermatrices comprising 276 nuc-proteins described above were summarized in Table 1.

**Table 1.** Supermatrices analyzed in this study.

| Supermatrix | Number of OTUs | Number of proteins<sup>1</sup> | Notes |
| --- | --- | --- | --- |
| nuc317 | 133 | 317 nuc-proteins | A supermatrix to infer the tree of global eukaryotes |
| nuc276 | 57 | 276 nuc-proteins | A supermatrix to infer the tree of Archaeplastida |
| nuc276ΔRP | 52 | 276 nuc-proteins | nuc276 without Picozoa or Rhodelphidia |
| nuc276ΔP | 54 | 276 nuc-proteins | nuc276 without Picozoa |
| nuc276ΔR | 55 | 276 nuc-proteins | nuc276 without Rhodelphidia |
| nuc276ΔP* | 54 | 276 nuc-proteins | Same as nuc276ΔP, but the average coverage of the rhodelphid sequences was lowered to 45.16% |
| nuc276ΔP** | 54 | 276 nuc-proteins | Same as nuc276ΔP, but the average coverage of the rhodelphid sequences was lowered to 49.99% |
| pld54 | 57 | 54 pld-proteins | A supermatrix to infer the tree of plastids |
| pld54Δg*r* | 51 | 54 pld-proteins | pld54 without six complex plastids |
| pld54Δr* | 53 | 54 pld-proteins | pld54 without red alga-derived complex plastids |
| pld54Δg* | 55 | 54 pld-proteins | pld54 without green alga-derived complex plastids |

<sup>1</sup> nuc-proteins, nucleus-encoded proteins; pld-proteins, plastid-encoded proteins.

#### Phylogenomic Analyses

We used IQ-TREE for phylogenetic analyses of nuc317 and nuc276 using the ML method (see below). The LG+C60+F+G model, accounting for both among-site rate variation approximated by gamma distributions (Yang and Kumer 1996) and the heterogeneity in amino acid frequency among alignment positions (Si Quang et al. 2008), was applied to tree searches and UFBoot2 (Hoang et al. 2018) that calculated ultrafast bootstrap values (UFBPs; 1,000 replicates). In non-parametric bootstrap analyses (100 replicates), the heterogeneity in amino acid frequency among alignment positions was approximated by the posterior mean site frequency model (Wang et al. 2018) estimated over the ML tree described above, coupled with the frequency vectors in the LG+C60+F+G model. nuc276 was also analyzed by Bayesian method by using PhyloBayes-MPI ver. 1.9 (Lartillot et al. 2013). We ran two Markov Chain Monte Carlo (MCMC) runs for 11,000 cycles with the GTR+CAT+G model (Lartillot and Philippe 2004) and stored trees every cycle during the MCMC runs.

Although the two MCMC runs did not converge (max-diff = 1), we discarded the trees from the initial 3,000 cycles, and the remaining trees were used to calculate the consensus tree and Bayesian posterior probabilities (BPPs). nuc276 and supermatrices derived from nuc276 (see below) were used for an approximately unbiased (AU) test (Shimodaira 2002), in which the number of RELL resampling was set to 10,000, to examine the relationship among Chloroplastida, Gaucophyta, and Rhodozoa. In the ML tree, Rhodozoa was grouped with the outgroup in the ML tree (Rhodo+Outg tree; see above). For the AU tests, we prepared two alternative trees of Archaeplastida, namely “Glauco+Outg tree” and “Chloro+Outg tree,” in which Glaucophyta and Chloroplastida were forced to be grouped with the outgroup, respectively. The two alternative trees were generated from the ML tree (i.e., Rhodo+Outg tree) by interchanging the three sub-clades manually.

Three modified versions of nuc276, “nuc276ΔPR,” “nuc276ΔR,” and “nuc276ΔP,” were prepared by excluding (i) two rhodelphids and three picozoan OTUs, (ii) the two rhodelphids, or (iii) the three picozoan OTUs from the original supermatrix, respectively (Table 1). Both ML and Bayesian analyses described above were repeated on nuc276ΔPR (the details of the analyses are the same as described above; in Bayesian analysis, the two MCMC runs did not converge as the max-diff stayed 1 after 11,000 cycles). In addition, the three supermatrices derived from nuc276 were subjected to the AU tests exploring the three candidate trees of Archaeplastida (see above). In each of the tests based on nuc276 and all of its derived supermatrices except for nuc276ΔRP, we compared the ML (Rhodo+Outg) tree inferred from a corresponding supermatrix and its manually modified trees (Chloro+Outg and Glauco+Outg trees) to each other. In contrast, we recovered the tree corresponding to the Glauco+Outg tree from the ML analysis of nuc276ΔRP (see below). Thus, for the test based on this supermatrix, the ML (Glauco+Outg) tree was compared with the two alternative trees corresponding to the Chloro+Outg and Rhodo+Outg trees that were generated by manual modification of the ML tree.

We evaluated the impact of fast-evolving positions in the original and modified versions of nuc276 on the ToA. nuc276 and its corresponding ML tree were subjected to IQ-TREE with the ‘--rate’ option to estimate the substitution rates for individual alignment positions (site-rates) using Bayesian approach. Then, the fastest-evolving positions were progressively removed from each of nuc276 and its derivatives at 20% intervals (fast-evolving position removal or FPR). We also removed random positions from each of the supermatrices described above at 20% intervals (random position removal or RPR). At each data point, 20 sets of alignment positions were chosen randomly and excluded from each of nuc276 and its derivatives to prepare 20 distinct RPR-processed supermatrices. This procedure allowed us to compare the phylogenetic inferences from a pair of nuc276ΔR and nuc276ΔP, from which the exact same set of randomly selected alignment positions was excluded (see above). The ML tree search, in conjunction with UFBoot2, was conducted on each of the supermatrices processed by both FPR and RPR.

The analyses of nuc276ΔR and nuc276ΔP revealed that rhodelphids and picozoan OTUs had different degrees of impact on the ToA (see above). We hypothesized that such a difference stemmed from the difference in coverage between the rhodelphid and picozoan sequences in the supermatrix. The average percent of missing data/gap positions in nuc276 was 4.85% in the rhodelphid sequences and 52.88% in the picozoan sequences. Thus, we artificially increased the proportions of missing data/gap positions in the rhodelphid sequences to the compatible level of those in the picozoan sequences (see below). First, we extracted 50,140 “empty positions” occupied by gaps in more than two picozoan sequences from nuc276. Likewise, we defined 44,767 of the positions in which gaps occurred less than two picozoan sequences in nuc276 as “filled positions” (note that “empty” and “filled” positions are complementary to each other in the picozoan sequences in nuc276). Then, we replaced any amino acid characters of the rhodelphid sequences at the positions corresponding to the “empty positions” in the picozoan sequences with gaps. This manipulation reduced the average percent value of missing/gap positions in the rhodelphid sequences to 54.84%, which is compatible with that in the picozoan sequences in the original nuc276 (Table 1). We then excluded the three picozoan OTUs from the modified nuc276 to generate “nuc276ΔP*”. We repeated the procedure described above, except for considering “filled positions” instead of “empty positions,” and generated “nuc276ΔP**” (the averaged percent value of missing/gap positions in the rhodelphid sequences to 50.01%; Table 1). Both nuc276ΔP* and nuc276ΔP** were used for the AU test assessing the three candidate trees of Archaeplastida, and the ML tree search coupled with UFBoot2 after FPR and RPR as described above.

### Plastid Protein-Based Phylogenomic Analyses

#### Supermatrices

We prepared a supermatrix based on pld-proteins by referring to that analyzed in Ponce-Toledo et al. (2017) to investigate the relationship among Chloroplastida, Glaucophyta, and Rhodophyta. From the NCBI RefSeq or GenBank database, we retrieved the protein (amino acid) sequences encoded in the plastid genomes of 42 archaeplastid species that were also considered in nuc276 (see above) plus six complex plastids that are the remnants of red/green algal endosymbionts, and nine cyanobacterial genomes that were chosen by referring to the cyanobacteria phylogeny in Strunecký et al. (2023) (57 OTUs in total). The 97 proteins encoded in the plastid genome of a glaucophyte *Cyanophora paradoxa* were set as queries, and their homologues were searched for in the protein sets encoded in the plastid and cyanobacterial genomes described above by blastp (options were set as ‘-evalue 1e-4 -max_target_seqs 5 -max_hsps 1’). In each blastp analysis, only the first hit was retained, as the proteins encoded in plastid genomes are usually single-copied or, even if duplicated, the copies are identical to each other at the amino acid level. We omitted 38 proteins that were absent from any of the chloroplastid species considered or contained only a single chloroplastid species, leaving 56 proteins for the analyses described below. The preliminary versions of 56 single-protein alignments were generated by MAFFT combined with trimAl for alignment trimming and then subjected to ML analyses using IQ-TREE for the final inspection. The ML trees inferred from two out of the 56 alignments were split into two subclades, both of which contained plastid sequences and were connected by a long internal branch. Such tree topologies resemble those containing paralogs (see above), and thus we decided to omit the two alignments bearing the suspicious phylogenetic ‘signal’ from the concatenation of the final supermatrix. The above-mentioned process yielded the final selection of 54 pld-proteins for the phylogenomic analyses described in the next section. These sequences were realigned by MAFFT with the L-INS-i algorithm, followed by a two-step trimming procedure composed of trimAl with ‘-gt 0.1’ option for the first step and BMGE with ‘-g 0.06 -e 0:0.5 -w 5’ options for the second step. The final versions of the 54 single-protein alignments were then concatenated into a “pld54” alignment (57 OTUs; 12,787 amino acid positions in total). We modified pld54 by excluding (i) the six complex plastids, (ii) two green alga-derived complex plastids, and (iii) four red alga-derived complex plastids to generate “pld54Δg*r*,” “pld54Δg*,” and “pld54Δr*,” respectively.

The supermatrices comprising 54 pld-proteins described above were summarized in Table 1. Note that, for both nuc276 and pld54, we sampled the archaeplastid species belonging to the same species or different but closely related species (those belonging to the same genera), except for two species belonging to Chloroplastida (*Pecluma dulcis* and *Lycopodium clavatum*) and three species belonging to Rhdophyta (*Rhodospora sordida, Ahnfeltia plicata*, and *Hildenbrandia rubra*) that were considered only in the latter supermatrix.

#### Phylogenomic Analyses

Both pld54 and pld54Δg*r* were subjected to a series of analyses using the ML methods, namely tree search, UFBoot2 (1,000 replicates), and non-parametric bootstrap analysis (100 replicates). Among the substitution models considering the heterogeneity in amino acid frequency among alignment positions by C60, the cpREV+C60+F+I+G model was selected by ModelFinder (Kalyaanamoorthy et al. 2017) implemented in IQ-TREE ver. 2.2.2.7. In non-parametric bootstrap analyses, the heterogeneity in amino acid frequency among alignment positions was approximated by the posterior mean site frequency model estimated over the ML tree described above, coupled with the frequency vectors in the cpREV+C60+F+I+G model. The two supermatrices were phylogenetically analyzed with Bayesian method using PhyloBayes-MPI. We ran two MCMC runs for 120,000 cycles with the GTR+CAT+G model and stored trees every cycle during the MCMC runs. The MCMC runs converged in the analysis of pld54 with a max-diff of 0.0903 and in the analysis of pld54Δg*r* with a max-diff of 0.0265. After discarding the trees from the initial 30,000 cycles as “burn-in,” the remaining trees were used to calculate the consensus tree and BPPs.

We conducted the pld54-based AU test comparing Chloro+Outg, Glauco+Outg, and Rhodo+Outg trees. Two alternative trees were prepared by modifying the ML tree inferred from the corresponding supermatrix (note that either Glauco+Outg or Chloro+Outg tree was recovered as the ML estimate, depending on the supermatrix analyzed and the phylogenetic method applied; see above). The site-lnLs were calculated with the cpREV+C60+F+I+G model over the ML and two alternative trees and used for the AU test (the number of RELL replicates was set to 10,000). We repeated the AU test considering the modified version of pld54 (i.e., pld54Δg*r*, pld54Δr*, and pld54Δg*; Table 1), together with the three test trees (see above). pld54 was modified by FPR and RPR, followed by the ML tree search and UFBoot2. The details were the same as described above, except for (i) the cpREV+C60+F+I+G model being applied to the site rate calculation and phylogenetic analyses, and (ii) removing alignment positions being repeated at 10% intervals. The above-mentioned set of analyses was repeated on pld54Δg*r*.

We additionally ran the ML tree search and UFBoot2 on pld54 after the exclusion of the positions bearing the bias in the amino acid composition identified by using WitChi (Köstlbacher et al. 2025, software version: git commit 3094f08). The program with the Wasserstein algorithm suggested removing 3,152 positions with amino acid composition bias from pld54, leaving 9,635 positions for the ML analysis.

We noticed the overall difference in branch length between the six complex plastids and others (i.e., primary plastids and cyanobacteria) in the pld54 phylogeny (see above). Thus, we focused on the alignment positions where the substitution rates were changed with and without the complex plastids and explored their impact on the phylogenetic inference of pld54. The substitution rate at each position (site-rate) based on pld54 was divided by the corresponding site-rate based on pld54Δg*r*. The alignment positions associated with site-rate ratios that departed largely from 1 potentially underwent “heterotachious” substitution models. A site-rate ratio >1 indicates that the site-rate up-shifts in the presence of the complex plastids relative to their absence. In the case of the site-rate down-shifting in the presence of the complex plastids relative to their absence, the ratio is supposed to be <1. We sorted the alignment positions in descending order of the site-rate ratio and progressively excluded the alignment positions associated with the 5% largest site-rate ratios and then 10-90% (at 10% intervals) from pld54. The resultant alignments were subjected to the ML tree search coupled with UFBoot2. We also excluded the alignment positions associated with the 5% smallest site-rate ratios, and then 10-90% progressively, followed by the ML tree search and UFBoot2 (see above). Lastly, we modified pld54 by excluding the alignment positions associated with both the 5% largest and 5% smallest site-rate ratios and subjected it to the ML tree search coupled with UFBoot2 and AU test examining the three types of ToA.

## Results

### Nuclear Protein-Based Phylogenomic Analyses

First, we phylogenetically analyzed nuc317 to examine whether the overall tree topology was changed before and after updating the samplings of taxa and proteins in a supermatrix. The ML phylogeny inferred from nuc317 (Fig. S1) verified the monophyly of Archaeplastida, as well as the sister relationship between Archaeplastida and Pancryptista (Yazaki et al. 2022). Other major clades of eukaryotes, such as TSAR, Haptista, Amorphea, CRuMs, Discoba, and Metamonada, were also recovered. The above-mentioned clades received UFBPs ranging from 88-100% and non-parametric ML bootstrap percent values (MLBPs) of 99-100% (Fig. S1). nuc276, a more compact alignment than nuc317, was generated for assessing the internal relationship among the three sub-clades in Archaeplastida. nuc276 comprised 45 archaeplastid species and the outgroup taxa (i.e., six pancryptists and six other eukaryotes). Any phototroph was omitted from the outgroup because such eukaryotes may have endosymbiotically transferred genes that can bias the ToA.

### Internal Relationship among the Three Sub-clades in Archaeplastida: With and Without Rhodelphidia and Picozoa

In the ML phylogeny of nuc276 (Fig. 2a), the monophyly of Archaeplastida was recovered. In the Archaeplastida clade, Chloroplastida and Glaucophyta are grouped together, excluding Rhodozoa. Thus, the overall ML tree corresponds to the Rhodo+Outg tree (Fig. 1c). The three bipartitions described above received full support from UFBoot2 and ML bootstrap (Fig. 2a; see also Fig. S2 for details). We provide the PhyloBayes tree inferred from nuc276 in Fig. S3 only for reference, as the two MCMC runs were not converged. We also examined the alternative tree topologies for the ToA by the AU test. In this test, the Rhodo+Outg tree was the ML estimate, and both Glauco+Outg and Chloro+Outg trees were rejected at the 1% level of statistical significance (Table 2).

**Table 2.** Comparisons of the three possible trees of Archaeplastida by AU tests based on nuc276 and its derivatives.

| Supermatrix | Sister to the outgroup | Difference in log likelihood<sup>1</sup> | p value<sup>2</sup> |
| --- | --- | --- | --- |
| nuc276 | **Rhodozoa** | ML (-4429234.956) | - |
|  | Glaucophyta | 131.18 | 0.00133** |
|  | Chloroplastida | 222.63 | 2.17e-11** |
| nuc276ΔRP | Rhodophyta | 39.887 | 0.156 |
|  | **Glaucophyta** | ML (-4098428.929) | - |
|  | Chloroplastida | 129.17 | 1.04e-05** |
| nuc276ΔP | **Rhodozoa** | ML (-4279548.599) | - |
|  | Glaucophyta | 71.633 | 0.0507 |
|  | Chloroplastida | 138.2 | 0.000105** |
| nuc276ΔR | **Rhodozoa** | ML (-4248542.293) | - |
|  | Glaucophyta | 31.496 | 0.235 |
|  | Chloroplastida | 178.79 | 3.28e-82** |
| nuc276ΔP* | **Rhodozoa** | ML (-4180765.582) | - |
|  | Glaucophyta | 22.239 | 0.284 |
|  | Chloroplastida | 120.18 | 6.22e-05** |
| nuc276ΔP** | **Rhodozoa** | ML (-4197249.135) | - |
|  | Glaucophyta | 20.598 | 0.316 |
|  | Chloroplastida | 110.72 | 0.000209** |

<sup>1</sup> For each ML tree, the corresponding log-likelihood value was given in parentheses.
<sup>2</sup> Double and single asterisks indicate that the AU test rejected the null hypothesis of no difference for the ML tree with $p < 0.01$ and $< 0.05$ , respectively.

**Figure 2.**
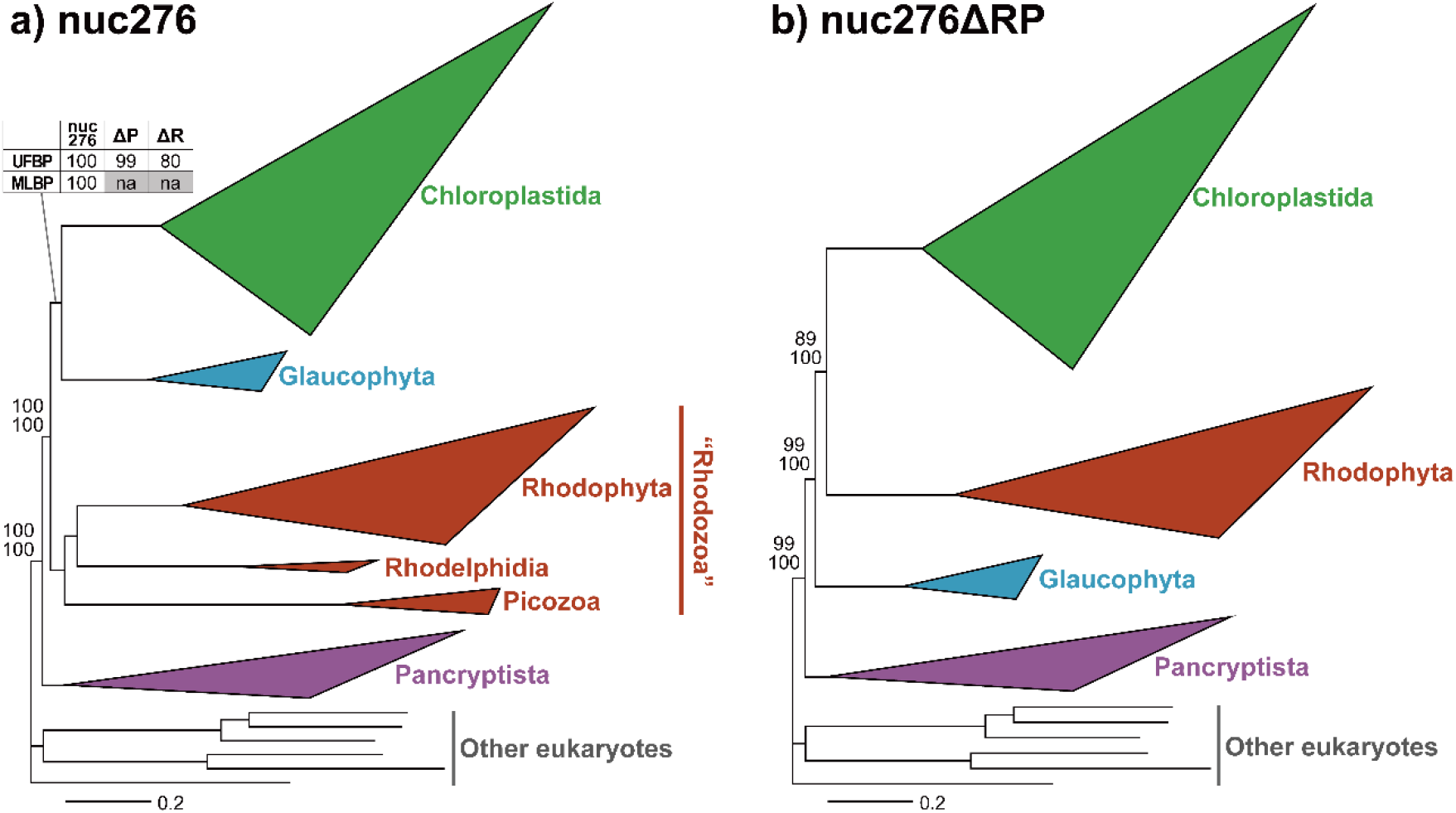
The trees of Archaeplastida inferred from nuc276 and nuc276ΔRP. The phylogenetic trees were recovered by the maximum likelihood (ML) method with the LG+C60+F+G model. The tree inferred from nuc276 and that from nuc276ΔRP are shown in (a) and (b), respectively. The sub-clades in Archaeplastida and Pancryptista were simplified into triangles. The size of each triangle is scaled to the number of the OTUs comprising the clade, and the longest and shortest distances from the ancestral to terminal nodes. All OTU names were omitted. We also omitted the support values for all bipartitions, except the values for (i) the CAM clade, (ii) the monophyly of Archaeplastida, and (iii) the union of Chloroplastida and Glaucophyta/Rhodophta (see Figs. S2 and S4 for the details). The support values shown on the top and bottom are ultrafast bootstrap percent values (UFBPs) and ML bootstrap values (MLBPs), respectively. The overall ML tree topologies inferred from nuc276ΔP and nuc276ΔR are essentially the same as that from nuc276 (see Fig. S7 for the details). The UFBPs for the bipartition uniting Chloroplastida and Glaucophyta calculated from nuc276ΔP and nuc276ΔR are listed in a table in (a).

The phylogenomic analyses presented in Yazaki et al. (2022) demonstrated that Rhodelphidia and *M. maris*, which split the branch leading to the Rhodophyta clade and that leading to the Cryptista clade, respectively, were critical to ameliorating the false affinity between Rhodophyta and Cryptista and helped to recover the Archaeplastida clade. When the analyses presented in Yazaki et al. (2022) were conducted, the transcriptome data of Picozoa reported in Schön et al. (2021) were not publicly available. Consequently, Yazaki et al. (2022) did not evaluate the impact of Picozoa on the ToA or its intimate phylogenetic affinity between Archaeplastida and Pancryptista. In this study, we explored the significance of Rhodelphidia and Picozoa to the relationship among the three sub-clades in Archaeplastida by analyzing the modified nuc276 excluding either or both of the non-photosynthetic relatives of Rhodophyta (i.e., nuc276ΔRP, nuc276ΔP, and nuc276ΔR; Table 1). In addition, we anticipated that the analyses of nuc276ΔRP provide clues for why the internal relationship in the ToA was incongruent between the phylogenomic trees based on nuc-proteins and pld-proteins, as no pld-protein data are available for the currently known members of Rhodelphidia or Picozoa that most likely lost their plastid genomes. Intriguingly, the Glauco+Outg tree was recovered from the ML analyses of nuc276ΔRP (Fig. 2b; see also Fig. S4 for the details). In the Archaeplastida clade received UFBP of 99% and MLBP of 100%, Chloroplastida and Rhodophyta were grouped, excluding Glaucophyta with UFBP of 89% and MLBP of 100% (Fig. 2b). As observed in Bayesian analysis of nuc276, the MCMC runs were not converged, and we hesitate to make any further discussion based on the PhyloBayes analysis of nuc276ΔRP (the PhyloBayes tree is provided in Fig. S5). The inference from nuc276ΔRP appeared not to be conclusive, as the AU test failed to reject the Rhodo+Outg tree with a *p*-value of 0.156 (Table 2).

### Analyses of Nuclear Protein Alignments Excluding Fast-Evolving and Random Positions

It is important to pursue why the exclusion of both Rhodelphidia and Picozoa switched the ML tree topology from the Rhodo+Outg to the Glauco+Outg tree (Figs. 1b and c). As Rhodelphidia and Picozoa split the somewhat long branch leading to the Rhodophyta clade (Fig. 2a), we wondered whether their absence triggered a phylogenetic artifact resulting in the position of Rhodophyta in the Archaeplastida clade instable. In general, fast-evolving positions in alignments are regarded as one of the typical sources of phylogenetic artifacts. Pioneering works have evaluated the degree of the contribution from fast-evolving positions to the ML tree by comparing the tree inferred from the original alignment and those from the alignments processed by FPR (Yazaki et al. 2022; Tikhonenkov et al. 2022; Eglit et al. 2024; Yazaki et al. 2025). We here subjected nuc276 and nuc276ΔRP to both FPR and RPR to examine the contribution of fast-evolving positions to the ML estimates from the two supermatrices.

The analyses of nuc276 processed by FPR and RPR suggested little contribution of fast-evolving positions in nuc276 to the dominance of the Rhodo+Outg tree over others. In Figures 3a-d, the results from FPR and RPR were presented as line graphs and box plots, respectively. The Rhodo+Outg tree was constantly recovered from the analyses after the removal of the top 20-60% fastest-evolving positions from nuc276 (Fig. 3b). After the top 80% fastest-evolving positions were removed, the remaining alignment positions were inappropriate to infer the relationship among the three sub-clades in the Archaeplastida clade, as Picozoa was tied to Glaucophyta instead of Rhodophyta and Rhodelphidia. In the analyses of RPR-processed alignments, the Rhodo+Outg tree was recovered at 20 out of the 20 trials (20-60% removals) and 14 out of the 20 trials (80% removal) (Fig. S6a). As the Rhodo+Outg tree was recovered in the analyses of both FPR and RPR-processed supermatrices, we conclude that the contribution of fast-evolving positions in nuc276 to the ML estimate (i.e., Rhodo+Outg tree) is negligible.

**Figure 3.**
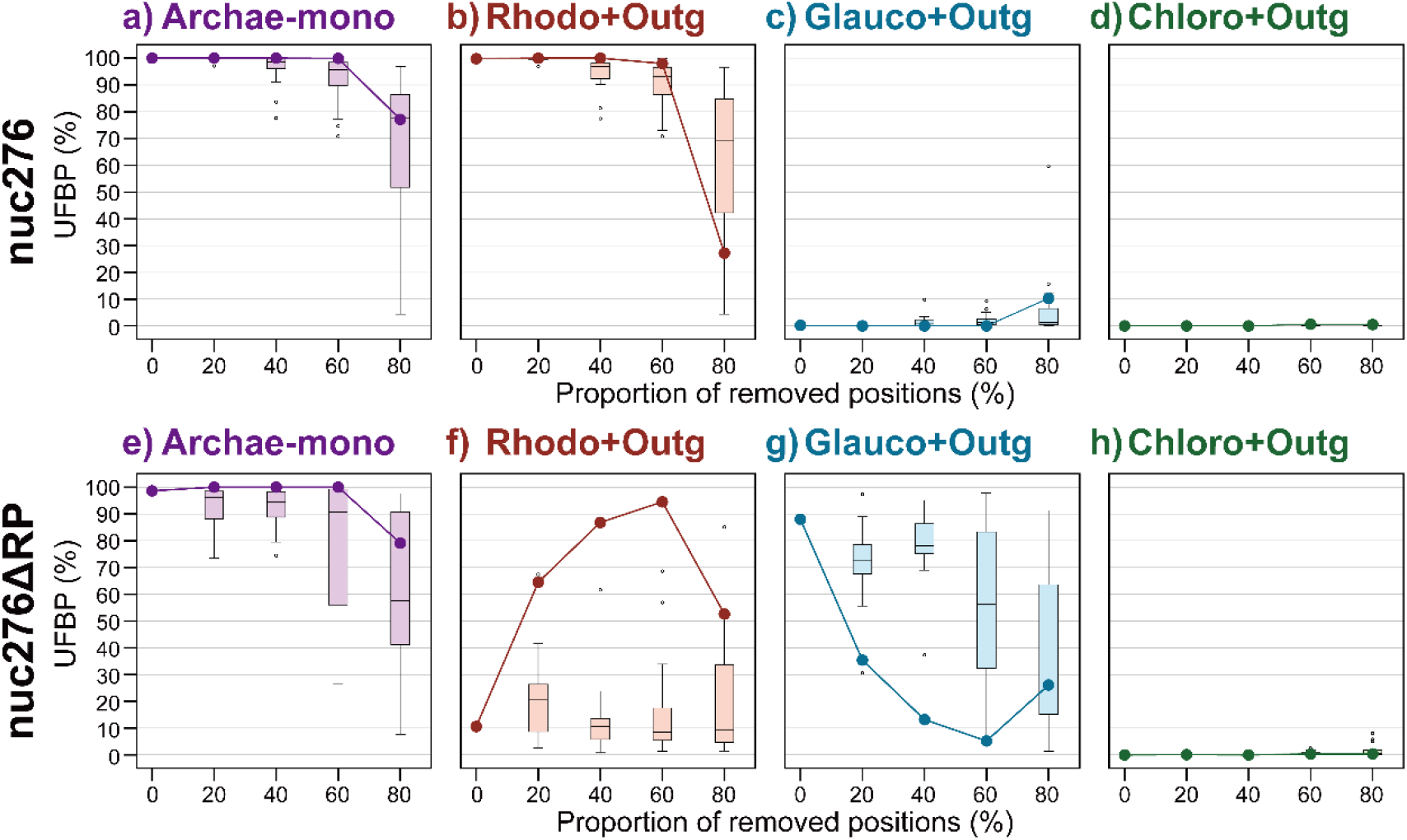
Analyses of nuc276 and nuc276ΔRP processed by the fast-evolving position removal (FPR) and random position removal (RPR) analyses. We removed the top 20%, 40%, 60%, and 80% of the fastest-evolving positions from nuc276 and nuc276ΔRP. The FPR-processed nuc276 and nuc276ΔRP were subjected to the ML tree search and UFBoot2 (1,000 replications). The ultrafast bootstrap percent values (UFBPs) for the monophyly of Archaeplastida (Archae-mono), the sister relationship between Rhodozoa/Rhodophyta and outgroup (Rhodo+Outg), that between Glaucophyta and outgroup (Glauco+Outg), and that between Chloroplastida and outgroup (Chloro+Outg) are plotted as line graphs in (a/e), (b/f), (c/g), and (d/h), respectively. In the ultrafast bootstrap analyses described here, we defined “Rhodo+Outg” trees satisfying both the monophyly of Archaeplastida and the separation of Rhodozoa/Rhodophyta from Glaucophyta and Chloroplastida. Likewise, “Glauco+Outg” trees satisfied the monophyly of Archaeplastida and the separation of Glaucophyta from Rhodozoa/Rhodophyta and Chloroplastida. We repeated the RPR-procedure for 20 times at each of four data points (i.e., 20, 40, 60, and 80% removals). Thus, 20 UFBPs were obtained at each data point and displayed using box plots.

We yielded the opposite conclusion from the analyses of nuc276ΔRP processed by FPR and RPR (Figs. 3e-h). The ML tree switched from the Glauco+Outg tree to the Rhodo+Outg tree after removing the top 20% fastest-evolving positions from nuc276ΔRP (see line graphs in Figs. 3f and g). The Rhodo+Outg tree appeared dominant over the Glauco+Outg tree until removing the top 80% fastest-evolving positions (Figs. 3f and g). In analyses excluding the top 80% fastest-evolving positions, the ML estimate was still the Rhodo+Outg tree, albeit the Glauco+Outg tree contended with the Rhodo+Outg tree according to UFBPs (the Rhodo+Outg and Glauco+Outg trees received UFBPs of 53% and 26%, respectively; see line graphs in Figs. 3f and g). In sharp contrast to the analyses of FPR-processed alignments described above, no switch of the ML tree topology from the Glauco+Outg tree to the Rhodo+Outg tree was promoted by the exclusion of random positions in nuc276ΔRP (see box plots in Figs. 3f and g). The Glauco+Outg tree was recovered in 12-19 out of the 20 trials in the analyses after removing 20-60% of random positions (Fig. S6b). We observed that the dominance of the Glauco+Outg tree over the Rhodo+Outg tree was diminished by further removal of random positions (Fig. S6b). The results from the analyses described here suggest that the Glauco+Outg tree relies heavily on the top 20% fastest-evolving positions, while the rest of the alignment positions favor the Rhodo+Outg tree rather than the Glauco+Outg tree.

The analyses of nuc276 processed by FPR and RPR suggested little contribution of fast-evolving positions in nuc276 to the dominance of the Rhodo+Outg tree over others. In Figures 3a-d, the results from FPR and RPR were presented as line graphs and box plots, respectively. The Rhodo+Outg tree was constantly recovered from the analyses after the removal of the top 20-60% fastest-evolving positions from nuc276 (Fig. 3b). After the top 80% fastest-evolving positions were removed, the remaining alignment positions were inappropriate to infer the relationship among the three sub-clades in the Archaeplastida clade, as Picozoa was tied to Glaucophyta instead of Rhodophyta and Rhodelphidia. In the analyses of RPR-processed alignments, the Rhodo+Outg tree was recovered at 20 out of the 20 trials (20-60% removals) and 14 out of the 20 trials (80% removal) (Fig. S6a). As the Rhodo+Outg tree was recovered in the analyses of both FPR and RPR-processed supermatrices, we conclude that the contribution of fast-evolving positions in nuc276 to the ML estimate (i.e., Rhodo+Outg tree) is negligible.

We yielded the opposite conclusion from the analyses of nuc276ΔRP processed by FPR and RPR (Figs. 3e-h). The ML tree switched from the Glauco+Outg tree to the Rhodo+Outg tree after removing the top 20% fastest-evolving positions from nuc276ΔRP (see line graphs in Figs. 3f and g). The Rhodo+Outg tree appeared dominant over the Glauco+Outg tree until removing the top 80% fastest-evolving positions (Figs. 3f and g). In analyses excluding the top 80% fastest-evolving positions, the ML estimate was still the Rhodo+Outg tree, albeit the Glauco+Outg tree contended with the Rhodo+Outg tree according to UFBPs (the Rhodo+Outg and Glauco+Outg trees received UFBPs of 53% and 26%, respectively; see line graphs in Figs. 3f and g). In sharp contrast to the analyses of FPR-processed alignments described above, no switch of the ML tree topology from the Glauco+Outg tree to the Rhodo+Outg tree was promoted by the exclusion of random positions in nuc276ΔRP (see box plots in Figs. 3f and g). The Glauco+Outg tree was recovered in 12-19 out of the 20 trials in the analyses after removing 20-60% of random positions (Fig. S6b). We observed that the dominance of the Glauco+Outg tree over the Rhodo+Outg tree was diminished by further removal of random positions (Fig. S6b). The results from the analyses described here suggest that the Glauco+Outg tree relies heavily on the top 20% fastest-evolving positions, while the rest of the alignment positions favor the Rhodo+Outg tree rather than the Glauco+Outg tree.

### Rhodelphidia and Picozoa as the Keys for Inferring the ToA

In the analyses of nuc276, we found no apparent evidence for the phylogenetic inference, in which Rhodozoa grouped with the outgroup, being severely biased (Figs. 3a-d). Notably, the results above reinforced the importance of Rhodelphidia and Picozoa for inferring the ToA, as the absence of the two lineages introduced the phylogenetic artifact stemming from fast-evolving positions (Figs. 3e-f). Yazaki et al. (2022) demonstrated that including Rhodelphidia (and M. maris) in phylogenomic analyses was critical to preventing the artifactual union of Rhodophyta and Cryptista, which was principally driven by cryptophytes, and recovering the monophyly of Archaeplastida. Note that the analyses of nuc276 and its derivatives contained no cryptophytes and, thus, were primarily immune to this phylogenetic artifact. Combining the results presented here with those reported in Yazaki et al. (2022), the non-photosynthetic lineages splitting from the branch leading to the Rhodophyta clade can be regarded as keys to inferring the early evolution of Archaeplastida.

Rhodelphidia and Picozoa were shown to suppress the phylogenetic artifact from fast-evolving positions. Although not described above, either of the two lineages alone suppressed the Glauco+Outg tree in the presence of the top 20% fastest-evolving positions. The Rhodo+Outg tree received UFBPs of 80-99% in the ML analyses of nuc276 modified by the reciprocal exclusion of Picozoa and Rhodelphidia (i.e., nuc276ΔP and nuc276ΔR; Figs. S7a and S7b). In the tests assessing the three types of the ToA based on nuc276ΔP and nuc276ΔR, the Rhodo+Outg tree received greater likelihoods than the Glauco+Outg or Chloro+Outg tree (Table 2). Nevertheless, the exclusion of Rhodelphidia alone and that of Picozoa alone appeared to bear different magnitudes in suppressing the Glauco+Outg tree. In the AU test considering Picozoa alone (i.e., nuc276ΔR), the Glauco+Outg tree received a large p value (0.235; Table 2), while the same tree barely failed to be rejected at the 5% level of statistical significance in the test considering Rhodelphidia alone (i.e., nuc276ΔP) (0.0507; Table 2). The results from the analyses of RPR-processed nuc276ΔP and nuc276ΔR are shown in Figures S6c, S6d, and S8a-d. The UFBPs for the Rhodo+Outg tree tend to be lower in the analyses considering Picozoa alone than in those considering Rhodelphidia as the sole basal lineage of Rhodophyta (ΔP and ΔR in Fig. S8b). In contrast, the UFBPs for the Glauco+Outg tree appeared to be elevated in the analyses considering Picozoa alone, compared to those considering Rhodelphidia alone (ΔP and ΔR in Fig. S8c). The dominance in the frequency of the recovery of the Rhodo+Outg tree after the removal of random positions, particularly in the removals of 60 and 80% random positions, was found to be loosened more in the absence of Rhodelphidia than in the absence of Picozoa, particularly (compare Figs. S6c and S6d).

We hypothesized that the difference in the proportion of gaps between the rhodelphid and picozoan sequences in nuc276 reflects the magnitude of suppressing the Glauco+Outg tree described above. To examine the hypothesis, nuc276ΔP* and nuc276ΔP** were prepared from nuc276 by excluding Picozoa and then increasing the gap proportion in the rhodelphid sequences to a similar degree to the picozoan sequences (Table 1). Importantly, the two distinct patterns of introducing gaps into the rhodelphid sequences produced nuc276ΔP* and nuc276ΔP**. Subsequently, the two supermatrices were subjected to the AU test considering the three types of the ToA and FPR/RPR followed by the ML tree search coupled with UFBoot2. The AU test based on nuc276ΔP* and nuc276ΔP** failed to reject the Glauco+Outg tree with large p values of 0.284 and 0.316 (Table 2); the overall dominance of the Rhodo+Outg tree over the Glauco+Outg tree based on the nuc276ΔP-derived supermatrices appeared to be more similar to that based on nuc276ΔR, rather than that based on nuc276ΔP (Table 2). Consequently, the increment of gap proportion in the rhodelphid sequences, regardless of how gaps being introduced, weakened the dominance of the Rhodo+Outg tree over the Glauco+Outg tree. A similar trend was observed in the analyses of RPR-processed nuc276ΔP* and nuc276ΔP** (Figs. S8a-d). Compared to the analyses of the RPR-processed nuc276ΔP, (i) the UFBPs for the Rhodo+Outg tree appeared to be lowered, while (ii) the UFBPs for the Glauco+Outg tree appeared to be elevated, in the analyses of the RPR-processed nuc276ΔP* and nuc276ΔP** (ΔP* and ΔP** in Figs. S8b and S8c). Regarding the ML tree topology, higher frequencies of recovering the Glauco+Outg tree were observed after the increment of gap proportion in the rhodelphid sequences (compare Figs. S6c and S6e, and S6c and S6f). Thus, we conclude that the difference in the magnitude of suppressing the Glauco+Outg tree between Rhodelphidia and Picozoa stemmed from the large proportion of gaps in the latter.

On the incongruity in the tree of Archaeplastida.

### Plastid Protein-Based Phylogenomic Analyses

#### Internal Relationship among the Three Sub-clades in Archaeplastida: With and Without Complex Plastids

As previously published phylogenetic studies of the supermatrices comprising multiple pld-proteins (Ponce-Toledo et al. 2017; Figueroa-Martinez et al. 2019), we recovered the Glauco+Outg tree from pld54 by the ML method with a UFBP of 80% and an MLBP of 67% (Fig. 4a; see Fig. S9 for details). On the other hand, Bayesian analysis of pld54 recovered the Chloro+Outg tree, demonstrating that the plastids in chloroplastids (green plastids) emerged prior to the separation of the plastids in rhodophytes and glaucophytes (BPP of 1.0; Fig. S11). In both ML and Bayesian analyses, the monophyly of plastids was recovered with UFBP of 100%, MLBP of 100%, and BPP of 1.0 (Fig. 4a). Corresponding with the uncertainties in the standard ML and Bayesian phylogenetic analyses (see above), the AU test comparing the Glauco+Outg tree (ML) with (i) the Chloro+Outg tree and (ii) the Rhodo+Outg tree assuming the emergence of the plastids in rhodophytes (red plastids) prior to the separation of glauco and green plastids failed to reject the Chloro +Outg tree (Table 3).

**Table 3.** Comparisons of the three possible trees of Archaeplastida by AU tests based on pld54 and its derivatives.

| Supermatrix | Sister to the outgroup<sup>1</sup> | Difference in log likelihood<sup>2</sup> | p value<sup>3</sup> |
| --- | --- | --- | --- |
| pld54 | red plastids+ | 36.703 | 0.000365** |
|  | **glauco plastids** | ML (-553602.8714) | - |
|  | green plastids+ | 0.70825 | 0.486 |
| pld54Δg*r* | red plastids | 38.506 | 3.09e-07** |
|  | glauco plastids | 3.631 | 0.424 |
|  | **green plastids** | ML (-489684.9682) | - |
| pld54Δg* | red plastids+ | 36.413 | 1.2e-05** |
|  | glauco plastids | 0.85283 | 0.477 |
|  | **green plastids** | ML (-529919.7079) | - |
| pld54Δr* | red plastids | 36.382 | 8.52e-06** |
|  | glauco plastids | 1.8319 | 0.461 |
|  | **green plastids+** | ML (-513445.9583) | - |
| pld54 after excluding the potential heterotachious positions | red plastids+ | 29.303 | 0.00351** |
|  | glauco plastids | 10.944 | 0.231 |
|  | **green plastids+** | ML (-513181.3828) | - |

<sup>1</sup> Plus signs indicate red or green alga-derived complex plastids were included.
<sup>2</sup> For each ML tree, the corresponding log-likelihood value is given in parentheses.
<sup>3</sup> Double and single asterisks indicate that the AU test rejected the null hypothesis of no difference for the ML tree with p < 0.01 and < 0.05, respectively.

**Figure 4.**
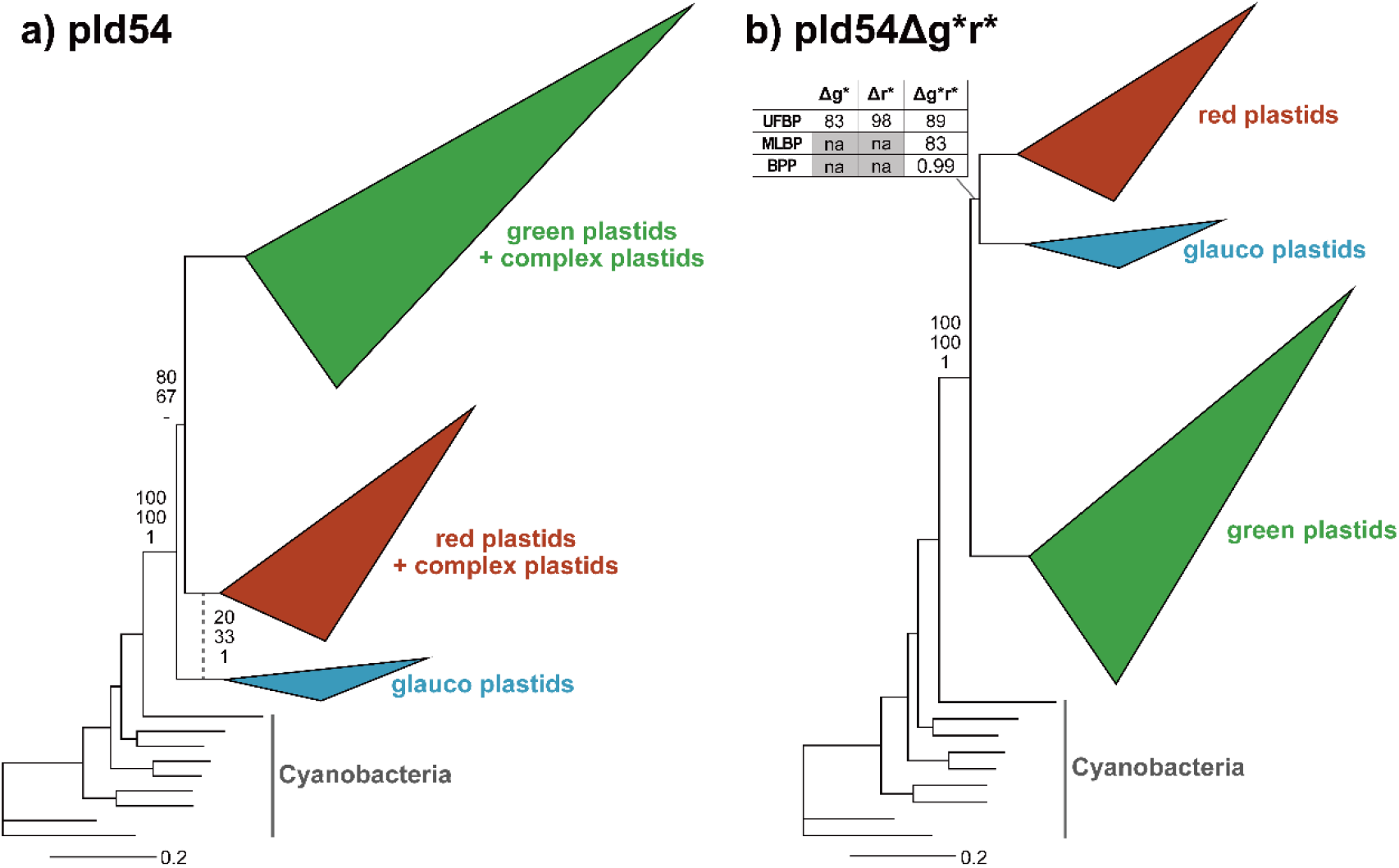
The trees of plastids inferred from pld54 and pld54Δg*r*. The phylogenetic trees were recovered by the maximum likelihood (ML) method with the cpREV+C60+F+I+G model. The tree inferred from pld54 and that from pld54Δg*r* are shown in (a) and (b), respectively. The three sub-clades in the plastid clade were simplified into triangles, as done in Fig. 2. We also omitted the support values for all bipartitions, except the support values for the monophyly of plastids and the sister relationship between two out of the three types of plastids. The support values shown on the top, middle, and bottom are ultrafast bootstrap percent values (UFBPs), ML bootstrap values (MLBP), and Bayesian posterior probabilities (BPPs), respectively. Bayesian analyses of pld54 recovered the union of the clade of red plastids and red alga-derived complex plastids and the clade of glauco plastids, which is represented by a dotted line in (a) (see Fig. S11 for the details). The statistical supports for the clade mentioned above are shown beside the dotted line. The overall ML tree topologies inferred from pld54Δg* and pld54Δr* are essentially the same as that from pld54Δg*r* (see Figs. S13 a & b). Thus, the UFBPs for the bipartition uniting red and glaucophyte plastids calculated from pld54Δg* and pld54Δr* are listed in a table in (b).

pld54 includes six complex plastids—two are the remnants of green algae engulfed by the ancestors of chlorarachniophytes and euglenids, and four represent the complex plastids, which can be traced back to red algal endosymbiont(s) directly or indirectly, in stramenopiles, a haptophyte alga, and a cryptophyte alga. The branches of the six complex plastids were longer than others in the plastid clade (Fig. S9), indicating that the substitution rates of their plastid-encoded genes were escalated during the reductive process that transformed algal endosymbionts into plastids in the host eukaryotic cells. We regard the six complex plastids in pld54 as dispensable for assessing the relationship among the three sub-clades in the plastid clade, as (i) they are essentially red or green plastids and (ii) their rapidly evolving natures may bias the phylogenetic inferences (e.g., potential sources of long-branch attraction artifacts). The ML and Bayesian analyses recovered the Chloro+Outg tree from pld54 after excluding the six complex plastids (pld54Δg*r*) with UFBP of 89%, MLBP of 83%, and BPP of 0.99 (Fig. 4b; see also Fig. S12 for the details), but the AU test remained the possibility for the Glauco+Outg tree (Table 3). Curiously, the ML analyses of pld54 excluding either green or red complex plastids (i.e., pld54Δg* or pld54Δr*) yielded the Chloro+Outg tree as well (Figs. S13a and S13b). The results described above clearly demonstrated that the recovery of the Glauco+Outg tree demands a pair of green and red complex plastids in a supermatrix.

#### Analyses of Plastid Protein Alignments Excluding Fast-Evolving and Random Positions

The phylogenetic relationship among the three sub-clades in the plastid clade inferred from pld54 appeared to be different with and without the six complex plastids (Figs. 4a and 4b). Thus, we subjected pld54 and pld54Δg*r* to FPR and RPR and subjected them to the ML tree search and UFBoot2 (Figs. 5a-h), as done on nuc276 and its derivatives. In the analyses of FPR-processed pld54, the monophyly of plastids was recovered until the removal of the top 80% fastest-evolving positions (see the line graph in Fig. 5a). The Glauco+Outg tree was dominant over the other two trees until the top 70% fastest-evolving positions were removed (see the line graphs in Figs. 5b-d), suggesting that fast-evolving positions are unlikely to be responsible for the position of glauco plastids relative to other two types of primary plastids in the plastid clade. In contrast, the Glauco+Outg and Chloro+Outg trees appeared to compete with each other in the analyses of RPR-processed pld54 (see the box plots in Figs. 5c and d). Regardless of the proportion of random positions removed, the UFBPs for the Glauco+Outg tree ranged widely from 0.1 to 99.7% and those for the Chloro+Outg tree from 0 to 99.9% (Figs. 5c and d). Similarly, the two trees were recovered from the RPR-processed pld54 at similar frequencies (Fig. S14a).

**Figure 5.**
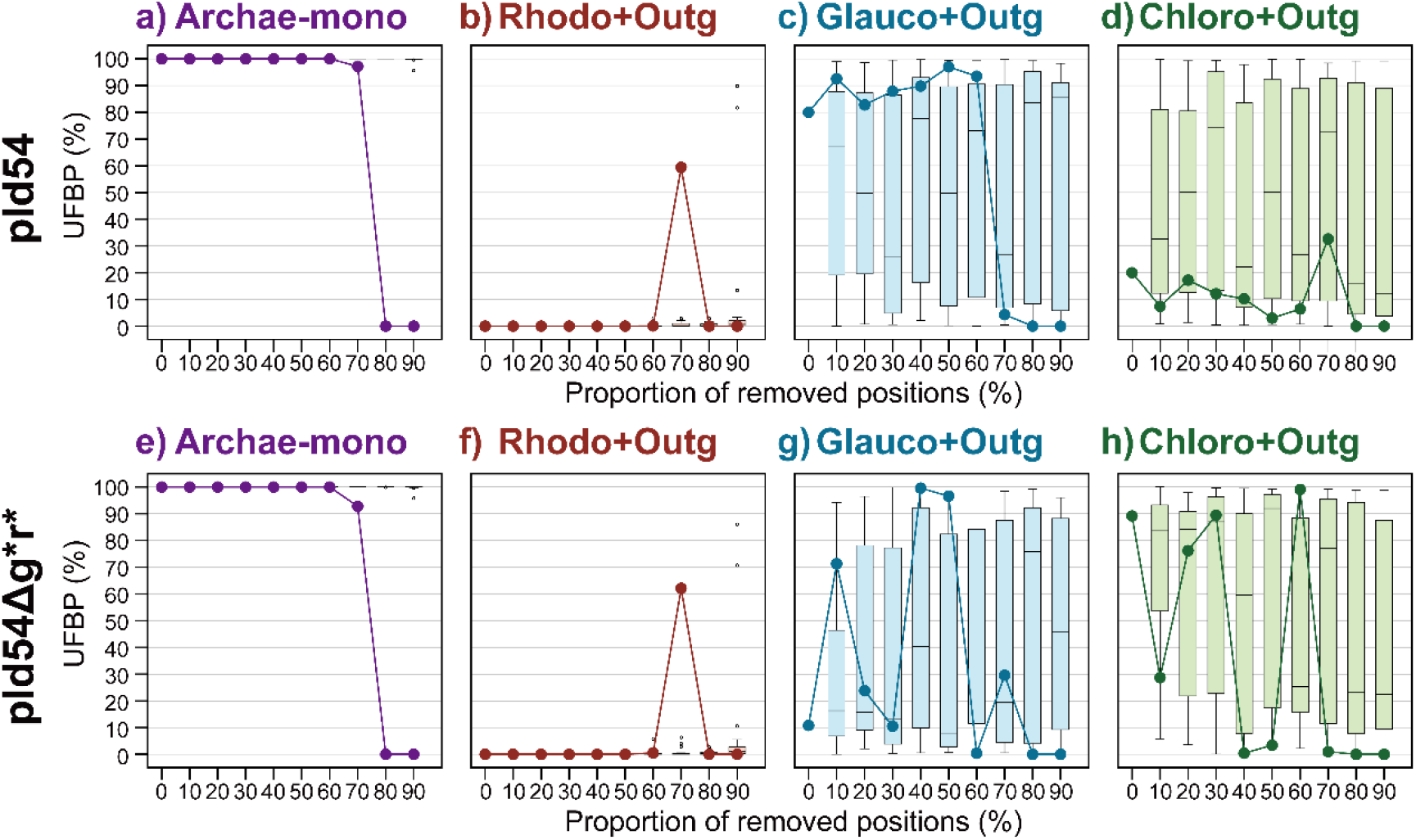
Analyses of pld54 and pld54Δg*r* processed by the fast-evolving position removal (FPR) and random position removal (RPR). We removed the top 10-90% of the fastest-evolving positions from pld54 and pld54Δg*r* at 10% intervals. The FPR-processed pld54 and pld54Δg*r* were subjected to the ML tree search and UFBoot2 (1,000 replications). The ultrafast bootstrap percent values (UFBPs) for the monophyly of plastids (Plastid-mono), red plastids with or without red alga-derived complex plastids grouped with the outgroup (Rhodo+Outg), glauco plastids grouped with the outgroup (Glauco+Outg), and green plastids with or without green alga-derived complex plastids grouped with the outgroup (Chloro+Outg) are plotted as line graphs in (a/e), (b/f), (c/g), and (d/h), respectively. In the ultrafast bootstrap analyses described here, we defined “Rhodo+Outg” trees satisfying both the monophyly of plastids and the separation of red plastids with or without red alga-derived complex plastids from the other plastids considered. Likewise, “Glauco+Outg” trees satisfied the monophyly of plastids and the separation of glauco plastids from the other plastids considered. We obtained 20 UFBPs calculated from 20 RPR-processed pld54/pld54Δg*r* at each of the nine data points (e.g., 10-90% removals) and present them in box plots.

**Figure 5.**
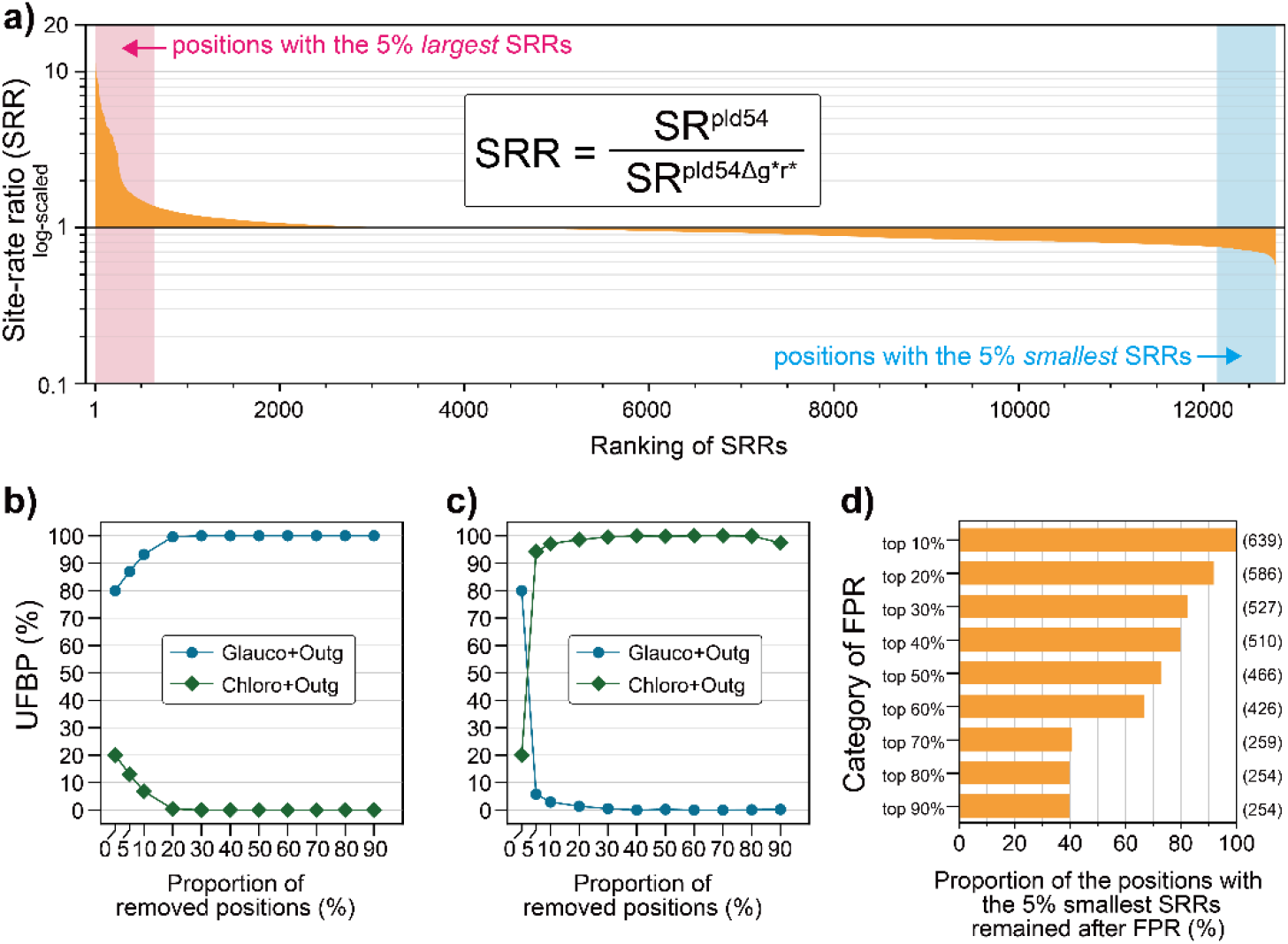
Analyses of pld54 and pld54Δg*r* processed by the fast-evolving position removal (FPR) and random position removal (RPR). We removed the top 10-90% of the fastest-evolving positions from pld54 and pld54Δg*r* at 10% intervals. The FPR-processed pld54 and pld54Δg*r* were subjected to the ML tree search and UFBoot2 (1,000 replications). The ultrafast bootstrap percent values (UFBPs) for the monophyly of plastids (Plastid-mono), red plastids with or without red alga-derived complex plastids grouped with the outgroup (Rhodo+Outg), glauco plastids grouped with the outgroup (Glauco+Outg), and green plastids with or without green alga-derived complex plastids grouped with the outgroup (Chloro+Outg) are plotted as line graphs in (a/e), (b/f), (c/g), and (d/h), respectively. In the ultrafast bootstrap analyses described here, we defined “Rhodo+Outg” trees satisfying both the monophyly of plastids and the separation of red plastids with or without red alga-derived complex plastids from the other plastids considered. Likewise, “Glauco+Outg” trees satisfied the monophyly of plastids and the separation of glauco plastids from the other plastids considered. We obtained 20 UFBPs calculated from 20 RPR-processed pld54/pld54Δg*r* at each of the nine data points (e.g., 10-90% removals) and present them in box plots.

The analyses of FPR-processed pld54Δg*r* recovered the monophyly of plastids until the top 80% fastest-evolving positions were removed (see the line graph in Fig. 5e), as observed in the analyses of the FPR-processed pld54 (Fig. 5a). Nevertheless, the ML estimate from an individual FPR-processed supermatrix appeared to be either Glauco+Outg or Chloro+Outg tree, depending on how far we removed fast-evolving positions from pld54Δg*r* (see the line graphs in Figs. 5g and h). The Glauco+Outg tree was preferred over the Chloro+Outg tree in the analyses after removing the top 10, 40, and 50% of the fastest-evolving positions. In contrast, the analyses of pld54Δg*r*, from which the top 20, 30, and 60% of the fastest-evolving positions were removed, recovered the Chloro+Outg tree rather than the Glauco+Outg tree. The contention between the Glauco+Outg and Chloro+Outg trees was reproduced in the analyses of RPR-processed pld54Δg*r*. As observed for RPR-processed pld54, the UFBPs for the two trees were distributed nearly from 0 to 100%, regardless of the proportion of alignment positions removed (see the box plots in Figs. 5g and h). Likewise, the two trees competed in terms of the frequency that occurred in the 20 trials at each of 10-70% removals of random positions (Fig. S14b).

The results described above provide us with two insights into the phylogenetic inferences from pld54 and pld54Δg*r*, namely (i) the limited phylogenetic signal to resolve the ToA (see below) and (ii) a type of systematic bias introduced by the six complex plastids to the phylogenetic inferences (see the next paragraph). First, the analyses of the RPR-processed pld54 demonstrated that the “preference” between the Glauco+Outg and the Chloro+Outg trees could be altered by choosing (even a slight amount of) alignment positions. For instance, we cannot be sure whether glauco or green plastids group with the outgroup, even after removing 10% of random positions from supermatrices (see the box plots in Figs. 5c and 5d). A similar “preference” between the Glauco+Outg and the Chloro+Outg trees was observed in the RPR-processed pld54Δg*r*. It is noteworthy that the analyses after the removal of 10-50% of random positions from pld54 tend to provide more support for the Glauco+Outg tree than the Chloro+Outg tree, as the UFBPs for the Glauco+Outg tree appeared to skew upward relative to the corresponding values from the analyses of RPR-processed pld54Δg*r* (compare Figs. 5c and 5g). Likewise, the Glauco+Outg tree was recovered more frequently in the analyses of RPR-processed pld54 than those of the RPR-processed pld54Δg*r* (Figs. S14a and S14b). Despite the upward skew of the UFBPs for the Glauco+Outg tree in the presence of the complex plastids hinting at a certain degree of the “signal” for the particular tree in the complex plastids, the analyses of the two supermatrices examined here are unlikely to clarify the internal branching pattern in the plastid clade with confidence.

Second, we here claim that the Glauco+Outg tree was most likely overrated by the complex plastids in the phylogenetic inferences from pld54. The ML estimate from pld54 was the Glauco+Outg tree (Figs. 4a and S9), albeit the same analysis excluding both or either green and red complex plastids yielded the Chloro+Outg tree (Figs. 4b, S12, and S13), suggesting that the union of green and red plastids relies primarily on the presence of both types of complex plastids in the supermatrix. Considering the evolutionary trajectories of green and red complex plastids and their rapidly evolving natures, they are dispensable to infer the relationship among green, glauco, and red plastids. Curiously, the putative overcredibility of the Glauco+Outg tree introduced by the two types of complex plastids has nothing to do with fast-evolving positions in pld54, as the analyses of the FPR-processed pld54 constantly recovered the Glauco+Outg tree (Fig. 5c). We can exclude the alignment positions biased in amino acid composition for the dominance of the Glauco+Outg tree. The overall tree topology remained the same before and after excluding the 3,152 positions bearing potential bias in amino acid composition (Fig. S10). Consequently, we then examine another aspect of sequence evolution, namely heterotachy (Lopez et al. 2002), for the reason why the Glauco+Outg tree is recovered from the phylogenetic analyses of pld54 in the next section.

#### Heterotachy is Responsible for the Union of Red and Green Plastids in the Analyses of Plastid Protein-Supermatrices

The alignment positions with the shifts in site-rate, which were introduced by the presence of the complex plastids, were identified based on the ratios of site-rates based on pld54 and those based on pld54Δg*r*. We calculated and sorted site-rate ratios (SRRs) by dividing the site-rates calculated based on pld54 by those calculated based on pld54Δg*r* as shown in Fig. 6a. In the alignment positions with SRRs > 1, the site-rates from the supermatrix including the complex plastids (i.e., pld54) are greater than the corresponding values from the same supermatrix including no complex plastid (i.e., pld54Δg*r*). The analyses of pld54 processed by the progressive removal of the alignment positions sorted by SRRs in descending order, regardless of the depth of the removal, recovered the Glauco-Outg tree as the ML estimate with UFBPs of 87-100% (Fig. 6b). Thus, we conclude that the alignment positions, in which the site-rates were accelerated by incorporating the complex plastids (i.e., SRR >1), are highly unlikely to be responsible for the dominance of the Glauco+Outg tree over the Chloro+Outg (or Rhodo+Outg) tree.

The removal of the alignment positions with the smallest SRRs (i.e., the site-rates calculated with the complex plastids were smaller than those without; see Fig. 6a) gave a drastic change in the ML estimate (Fig. 6c). The removal of the positions with the 5% smallest SRRs, which correspond to those shaded in blue in Fig. 6a, switched the ML estimate from the Glauco+Outg tree to the Chloro+Outg tree (Fig. 5c). Throughout this set of the removal of the positions based on the SRR, the Chloro+Outg tree was constantly recovered as the ML estimates with UFBPs of 94.2-100%. In sharp contrast, the UFBPs for the Glauco+Outg tree were negligible (0-5.8%). The results described above claim that the “signal” for the Glauco+Outg tree is concentrated in the alignment positions with the 5% smallest SRRs. In other words, the Glauco+Outg tree principally depends on the heterotachy introduced by the complex plastids. Importantly, our claim above can reconcile the results from the analyses of the FPR-processed pld54 (Figs. 5c). Even after the removal of the top 60% fastest-evolving positions from pld54, almost two-thirds of the alignment positions with the 5% smallest SRRs remained for tree search and UFBoot2 (Fig. 6d), resulting in the recovery of the Glauco+Outg tree as the ML estimate.

We reconstructed the ToA from pld54 after excluding the alignment positions with heterotachy. To be fair, the ML analysis was conducted on pld54 after excluding the alignment positions associated with the largest 5% site-rate ratios, of which exclusion yielded no change in the ML estimate (Fig. 6b), as well as those with the smallest 5% site-rate ratios (i.e., the areas shaded in pink and blue in Fig. 6a). As anticipated from the analyses of pld54 processed by removing the alignment positions with the smallest 5% site-rate ratios (Fig. 6c), the Chloro+Outg tree was recovered as the ML estimate with a UFBP of 90% (Fig. S15), albeit the AU test failed to reject the Glauco+Outg tree (*p* = 0.231; Table 3). Combined with the results derived from the analyses of the RPR-processed pld54 and pld54Δg*r* (see the previous section), we again conclude that pld54 retains no sufficient signal to clarify the internal branching pattern in the plastid clade with confidence.

### Can Analyses with Complex Models Resolve the Incongruity Between Nuc-protein- and Pld-protein-Based ToA?

The systematic assessments on supermatrices of nuc-proteins (nuc-supermatrices; nuc276 and its derivatives) and those of pld-proteins (pld-supermatrices; pld54 and its derivatives) demonstrated that, depending on taxon sampling, both types of the phylogenomic inference of the ToA were severely biased for distinct reasons. Nevertheless, we have so far failed to resolve the incongruity between the ToA inferred from the nuc-supermatrix and that from the pld-supermatrix. Currently, there is no practical procedure to pursue the issue in the ToA addressed here, except for applying more complex substitution models than those used in the analyses mentioned above (i.e., LG+C60+F+G and cpREV+C60+F+I+G) (Table 4). We additionally examined two complex models, namely a model in which among-site rate heterogeneity was approximated by a FreeRate model (LG+C60+F+R9; Soubrier et al. 2012), and GHOST models (Crotty et al. 2020) accounting for heterotachy in a supermatrix (LG+C60+F+H4 and LG+C60+F+H8), in the AU tests comparing the Chloro+Outg, Glauco+Outg, and Rhodo+Outg trees over nuc276. nuc276 was chosen because no apparent phylogenetic bias was detected (see above). Due to the severe computational burdens of the FreeRate and GHOST models, these models are not practical for the ML tree search, UFBoot2, or non-parametric bootstrap. Importantly, none of the three complex models brought any substantial change in the results of the AU tests, dominating the Rhodo+Outg tree over the alternatives (Table 4).

**Table 4.** Comparisons of the three possible trees of Archaeplastida by AU tests with the FreeRate and GHOST models.

| Alignment | Substitution model (BIC<sup>1</sup>) | Sister to the outgroup<sup>2</sup> | Difference in log likelihood<sup>3</sup> | p value<sup>4</sup> |
| --- | --- | --- | --- | --- |
| nuc276 | LG+C60+F+G<br>(8,860,658.897) | **Rhodozoa** | ML (-4429234.956) | - |
|  |  | Glaucomphyta | 131.18 | 0.00133** |
|  |  | Chloroplastida | 222.63 | 2.17e-11** |
|  | LG+C60+F+R9<br>(8,843,760.388) | **Rhodozoa** | ML (-4420699.747) | - |
|  |  | Glaucomphyta | 113.18 | 0.0026** |
|  |  | Chloroplastida | 196.28 | 1.29e-87** |
|  | LG+C60+F+H4<br>(8,850,693.831) | **Rhodozoa** | ML (-4422332.764) | - |
|  |  | Glaucomphyta | 140.01 | 0.000224** |
|  |  | Chloroplastida | 226.71 | 2.55e-06** |
|  | LG+C60+F+H8<br>(8,823,860.076) | **Rhodozoa** | ML (-4406348.7) | - |
|  |  | Glaucomphyta | 89.708 | 0.0318* |
|  |  | Chloroplastida | 175.88 | 1.64e-50** |
| pld54 | cpREV+C60+F+I+G<br>(1,109,021.330) | red plastids+ | 36.703 | 0.000365** |
|  |  | **glauco plastids** | ML (-553602.8714) | - |
|  |  | green plastids+ | 0.70825 | 0.486 |
|  | cpREV+C60+F+I+R7<br>(1,106,108.846) | red plastids+ | 37.529 | 2.45e-05** |
|  |  | **glauco plastids** | ML (-552094.6204) | - |
|  |  | green plastids+ | 1.3258 | 0.47 |
|  | cpREV+C60+F+I+H4<br>(1,105,792.557) | red plastids+ | 37.785 | 1.35e-05** |
|  |  | **glauco plastids** | ML (-550404.574) | - |
|  |  | green plastids+ | 3.48 | 0.439 |
|  | cpREV+C60+F+I+H8<br>(1,102,983.706) | **red plastids+** | ML (-546860.8501) | - |
|  |  | glauco plastids | 21.113 | 0.231 |
|  |  | green plastids+ | 60.204 | 0.0123* |

<sup>1</sup> Bayesian information criterion values were calculated from the log-likelihoods of the Rhodo+Outg tree inferred from nuc276, or that of the Glauco+Outg tree inferred from pld54.
<sup>2</sup> Plus signs indicate the inclusion of red or green complex plastids.
<sup>3</sup> For each ML tree, the corresponding log-likelihood value is given in parentheses.
<sup>4</sup> Double and single asterisks indicate that the AU test rejected the null hypothesis of no difference for the ML tree with $p < 0.01$ and $< 0.05$ , respectively.

For the pld-supermatrix for the AU tests with a FreeRate model (cpREV+C60+F+I+R7) and GHOST models (cpREV+C60+F+I+H4 and cpREV+C60+F+I+H8), we selected pld54. As discussed above, pld54 is most likely short of the phylogenetic signal to resolve the ToA. Rather, this series of the AU test was conducted to examine whether the GHOST model can overcome the recovery of the Glauco+Outg tree depending solely on the heterotachious positions in the supermatrix. The overall results (i.e., the rejection of the Rhodo+Outg tree) were unchanged by switching the approximation of among-site rate heterogeneity from the gamma to the FreeRate model (Table 4). Likewise, the test applying the cpREV+C60+F+I+H4 model gave the highest log-likelihood to the Glauco+Outg tree and rejected the Rhodo+Outg tree at the 1% level of statistical significance (Table 4). Intriguingly, a drastic change occurred in the result from the test incorporating a more parameter-rich model (cpREV+C60+F+I+H8); the Rhodo+Outg tree received a higher log-likelihood than the Glauco+Outg or Chloro+Outg tree (Table 4). It is attractive to reconcile that not the H4 but the H8 model ameliorated sufficiently the heterotachy in pld54, inferring the Rhodo+Outg tree from pld54. There is no evident sign of overparameterization in the H8 model. So far, the H8 model was judged to be more appropriate than the other models examined here under the Bayesian information criterion (Table 4). The result described above is encouraging, but we need to better understand why the two GHOST models (i.e., H4 and H8 models) yielded two drastically different results in the AU tests based on pld54 in the future.

## Discussion

We detected no apparent sign of methodological bias in the phylogenetic inferences from nuc276 as long as Rhodelphidia and/or Picozoa are present in the supermatrix (see the first section in the Results). On the other hand, the analyses of pld54 demonstrated that this supermatrix is fundamentally short to resolve the relationship among red, glauco, and green plastids, but bears the heterotachy introduced by the complex plastids, which overrates the phylogenetic affinity between red and green plastids (see the second section in the Results). We recommend excluding complex plastids, for which the tempo and mode of substitutions are substantially different from those of primary plastids or cyanobacteria, from future studies assessing the ToA based on pld-supermatrices. Alternatively, the putative false impact of the complex plastids on the ToA can be accommodated by applying a more realistic substitution model than those mainly applied in this study, e.g., one that takes into account heterotachy, for future phylogenetic analyses of pld-supermatrices (see the third section in the Results). Based on the results from the phylogenetic analyses of nuc276 and pld54, we favor the Rhodo+Outg tree as the working hypothesis on the ToA for future examinations. Apparently, this conclusion is tentative and should be examined rigorously in future studies. In this section, we evaluate potential approaches toward resolving the true ToA in the future.

One can argue that the issue of insufficient resolution in pld54 can be overcome by preparing and analyzing new pld-supermatrices larger than pld54 in size. There is a large difference in size between nuc276 and pld54 (i.e., 94,907 and 12,787 amino acid positions), and such a size difference likely resulted in the uncertainty remaining in the phylogenetic inference from pld54. In theory, we can generate a larger supermatrix by combining the proteins encoded in plastid genomes and nucleus-encoded plastid proteins in archaeplastid species. We do not exclude such ideas, but need to be extremely cautious about which proteins to include, as they must have been inherited vertically from the cyanobacterial endosymbiont without any replacement by a distantly related homologue.

Besides phylogenetic analyses of pld-proteins and nuc-proteins, the ToA can be inferred from the proteins encoded in mitochondrial genomes (mt-proteins). In reality, alignments comprising multiple mt-proteins (mt-supermatrices) may suffer from the issue of size, as seen in pld-supermatrices (see above). Because of the number of mt-proteins being restricted, the size of a mt-supermatrix cannot be compatible with that of nuc276 or even smaller than pld54. For instance, some of us prepared and analyzed an alignment of 11 mt-proteins (3,533 amino acid positions in total) to assess whether *Palpitomonas bilix*, a kathablepharid, goniomonads, and cryptophyte algae formed a clade (Nishimura et al. 2020). Both ML and Bayesian trees recovered the monophyly of the species of interest (i.e., cryptists) but failed to group Chloroplastida, Glaucophyta, and Rhodophyta into a single clade. The phylogenetic inference from a similar mt-supermatrix (14 mt-proteins; 3,267 amino acid positions) recovered the monophyly of Archaeplastida, but the statistical support from the ML method was poor (Jackson and Reyes-Prieto 2014). Thus, mt-supermatrices are highly unlikely to be appropriate for inferring the ToA with confidence.

In the end, we contemplate that novel members of Archaeplastida, if they exist, may play a pivotal role in inferring the ToA. Such novel species are not entirely speculative, as both Rhodelphidia and Picozoa were recently documented as novel members in Archaeplastida (Gawryluk et al. 2019; Schön et al. 2021). Thus, novel members, which are photosynthetic and branch at the base of Chloroplastida, Glaucophyta, or Rhodophyta, may have been overlooked in natural environments. For instance, suppose that the Rhodo+Outg tree of Archaeplastida is genuine, and the analyses of pld-supermatrices conducted to date falsely disfavored the ‘true’ tree. The pld-protein data of photosynthetic species currently unknown to science may change the relationship among red, glauco, and green plastids drastically or at least help the Rhodo+Outg tree compete with the other two trees in the analyses of pld-supermatrices. On the contrary, there still remains the possibility of the analyses of nuc-supermatrices being biased by unknown reasons, and thus, the Glauco+Outg or Chloro+Outg tree can be the genuine ToA. If so, as-yet-unknown species, particularly those branched at the base of Chloroplastida or Glaucophyta, may suppress the dominance of the Rhodo+Outg tree in the analyses of the current nuc-supermatrices.

## Conclusion

The internal relationship among the subclades in Archaeplastida appeared not to be as straightforward as initially anticipated, as our analyses demonstrated that the phylogenetic inferences from both nuc- and pld-supermatrices suffer from distinct systematic biases depending on taxon sampling. We finally propose the Rhodo+Outg tree as a working hypothesis for the ToA, but there is a large room for further validation on this matter. Fortunately, a part of our analyses presented above hinted that complex substitution models—for instance, those incorporating heterotachy—are on track to be pursued in future analyses of pld-supermatrices to unveil the true ToA. It is also important to find and incorporate novel members of Archaeplastida, particularly those branching at the base of the three subclades, into future studies, as they may be of great help in recovering the true ToA. Nevertheless, we might not resolve the possibility of the incongruity between the ToA inferred from a nuc-supermatrix and that from a pld-supermatrix remaining even after the improvements of the methodology in tree reconstruction and taxon sampling in supermatrices. In that case, we need to explore biological reasons that can reconcile the incongruity challenged in this study. For instance, we may evaluate the potential impact of incomplete lineage sorting or lateral gene transfer on nuclear genome evolution in Archaeplastida. Depending on the situation, it might also be necessary to reconsider a single origin of primary plastids in Chloroplastida, Glaucophyta, and Rhodophyta.

## Acknowledgements

We thank Dr. Takeshi Nakayama (University of Tsukuba, Japan) for providing the light microscopic images used in Fig. 1a. R. H. thanks Dr. Andrew J. Roger (Dalhousie University, Canada) for helpful advice and discussion. Computations were partially performed on the NIG supercomputer at ROIS National Institute of Genetics. This work was supported by grants from the Japan Society for the Promotion of Science (JSPS) (23K27226 and BPI06050 awarded to Y. I., and 24K09587 to T. N.). R. H. was supported by the JSPS Overseas Research Fellowship. R. I. was supported by JST SPRING, Grant number JPMJSP2124.

## Data availability

Supplementary data are available at https://doi.org/10.5061/dryad.b2rbnzssh.

