## Supplementary figures for "Phylogenetic Dissection Provides Insights into the Incongruity in the Tree of Archaeplastida Between the Analyses of Nucleus- and Plastid-Encoded Proteins"

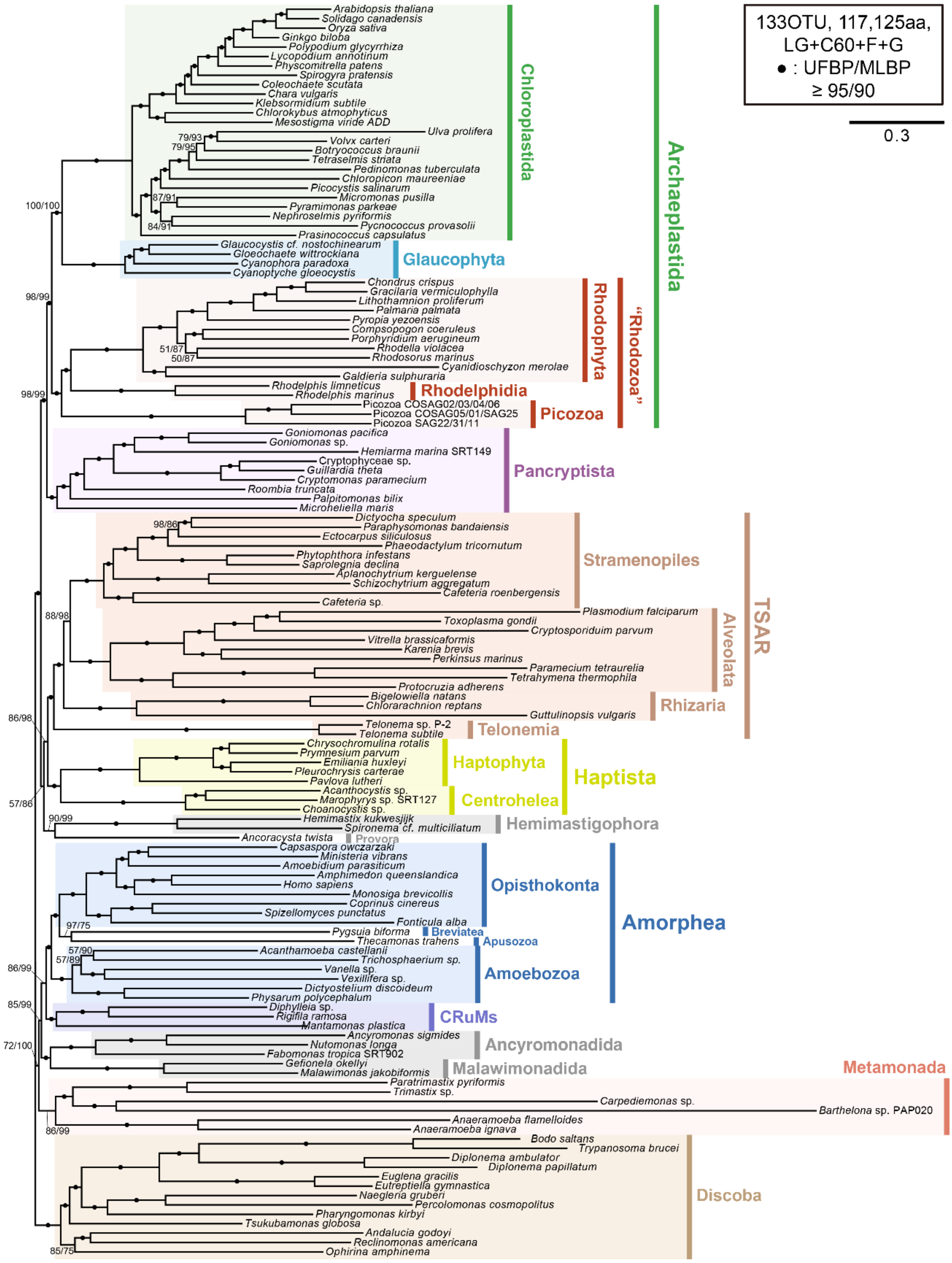


**Supplementary figure 1. The global eukaryotic phylogeny inferred from nuc317.** The phylogenetic tree was recovered by the maximum likelihood (ML) method. For each bipartition, an ultrafast bootstrap percent value (UFBP) and ML bootstrap values (MLBPs) are shown on the left and right, respectively. The bipartitions marked by dots received UFBPs ≧95% and MLBPs ≧90%.

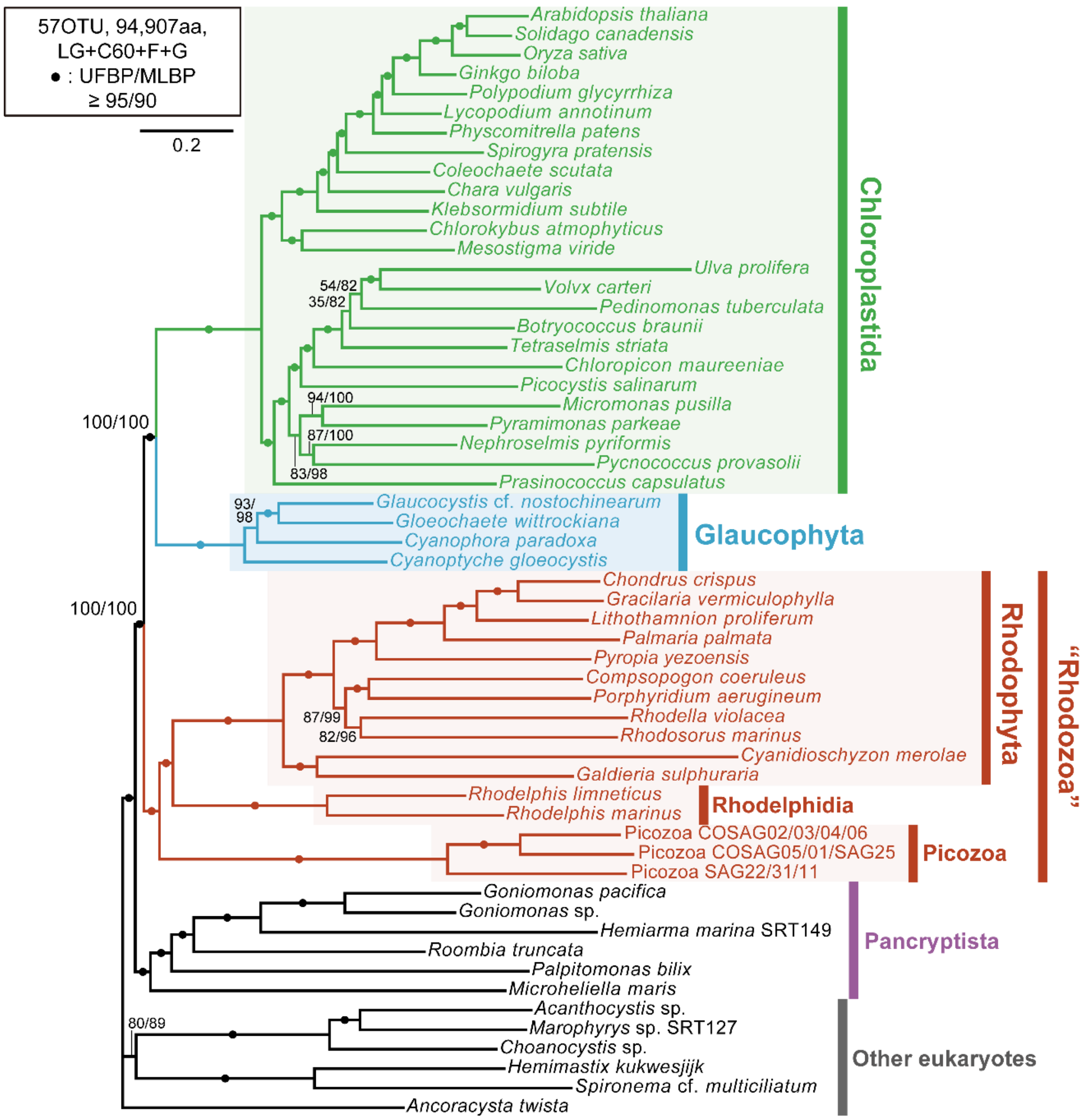


**Supplementary figure 2. The maximum likelihood tree of Archaeplastida inferred from nuc276.** The phylogenetic tree was recovered by the maximum likelihood (ML) method. For each bipartition, an ultrafast bootstrap percent value (UFBP) and ML bootstrap values (MLBPs) are shown on the left and right, respectively. The bipartitions marked by dots received UFBPs ≧95% and MLBPs ≧90%.

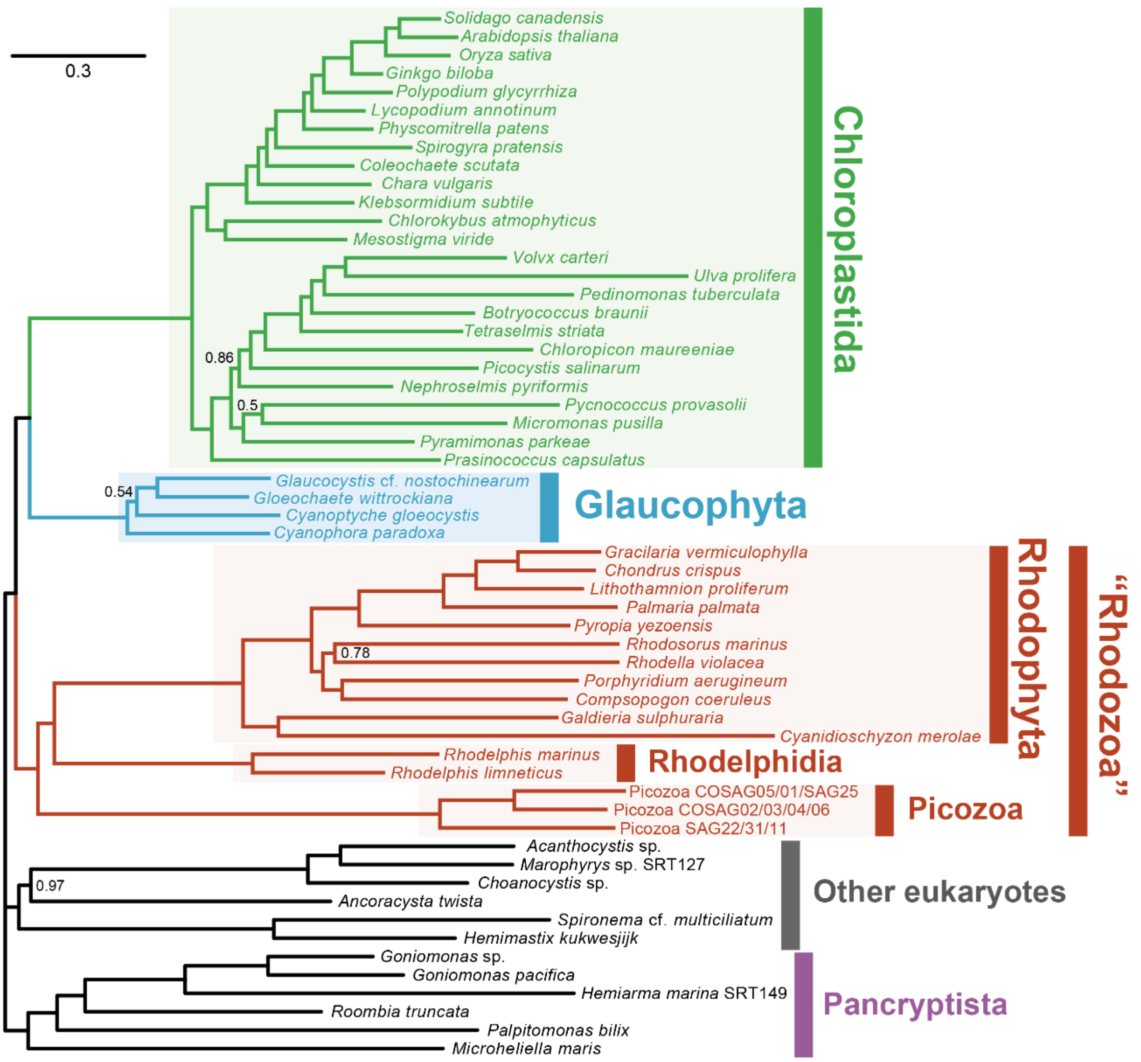


**Supplementary figure 3. Bayesian tree of Archaeplastida inferred from nuc276.** The phylogenetic tree was recovered by Bayesian method. Note that the two MCMC chains were not converged. All the bipartitions received Bayesian posterior probabilities of 1.0, except for those with the values specified.

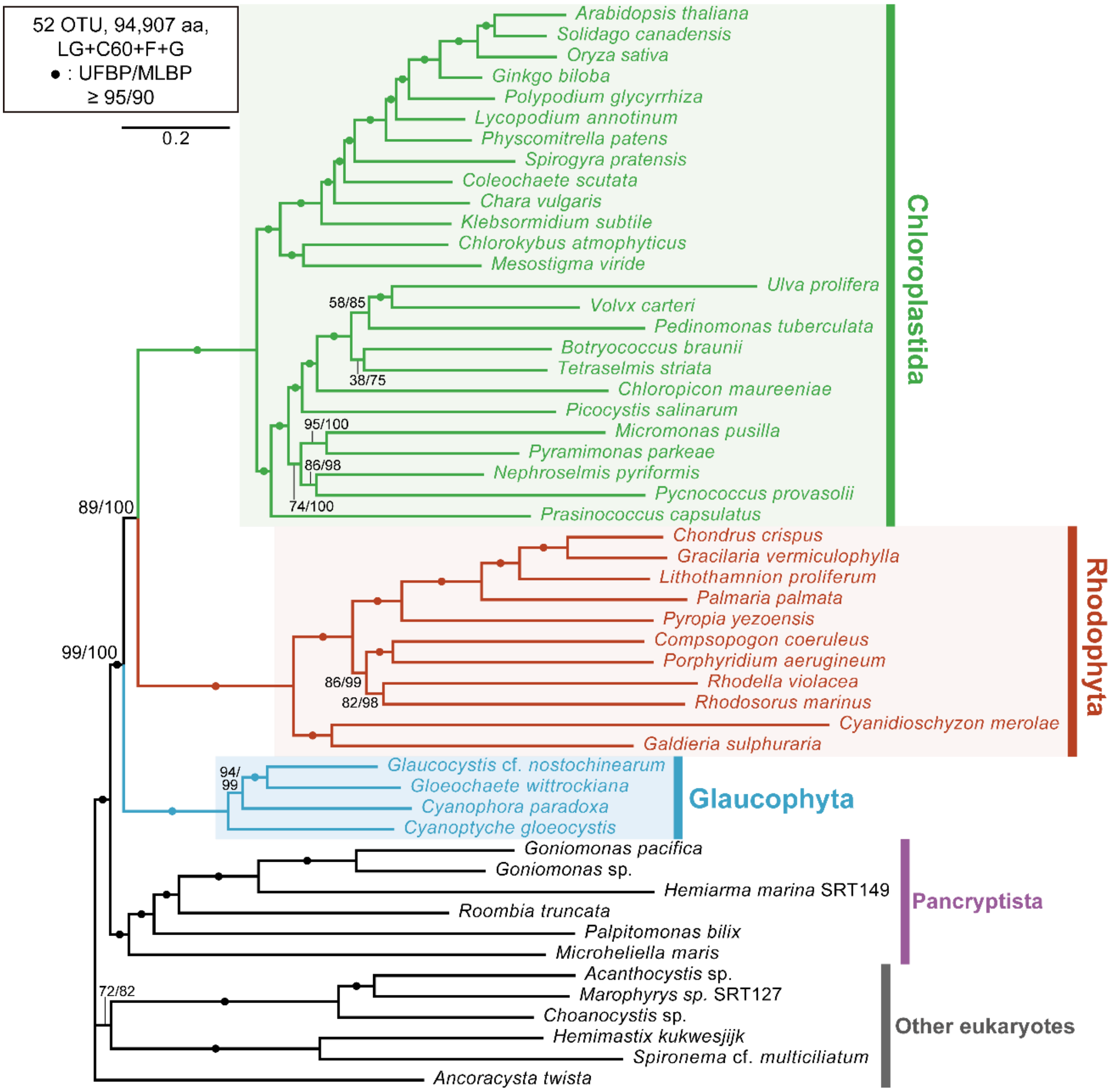


**Supplementary figure 4. The maximum likelihood tree of Archaeplastida inferred from nuc276ΔRP.** The details of this figure are the same as those of supplementary figure 2.

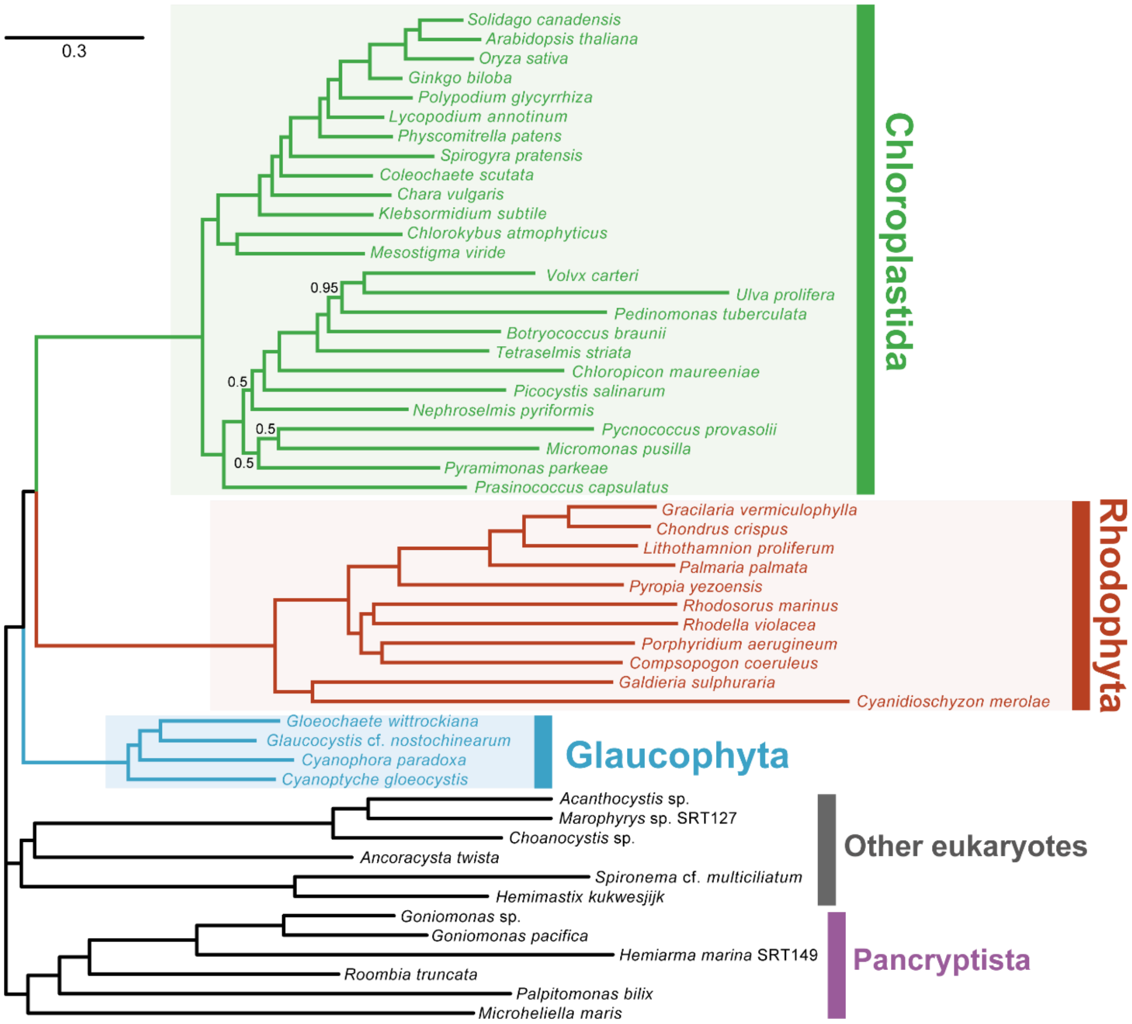


**Supplementary figure 5. Bayesian tree of Archaeplastida inferred from nuc276ΔRP.** The details of this figure are the same as those of supplementary figure 3.

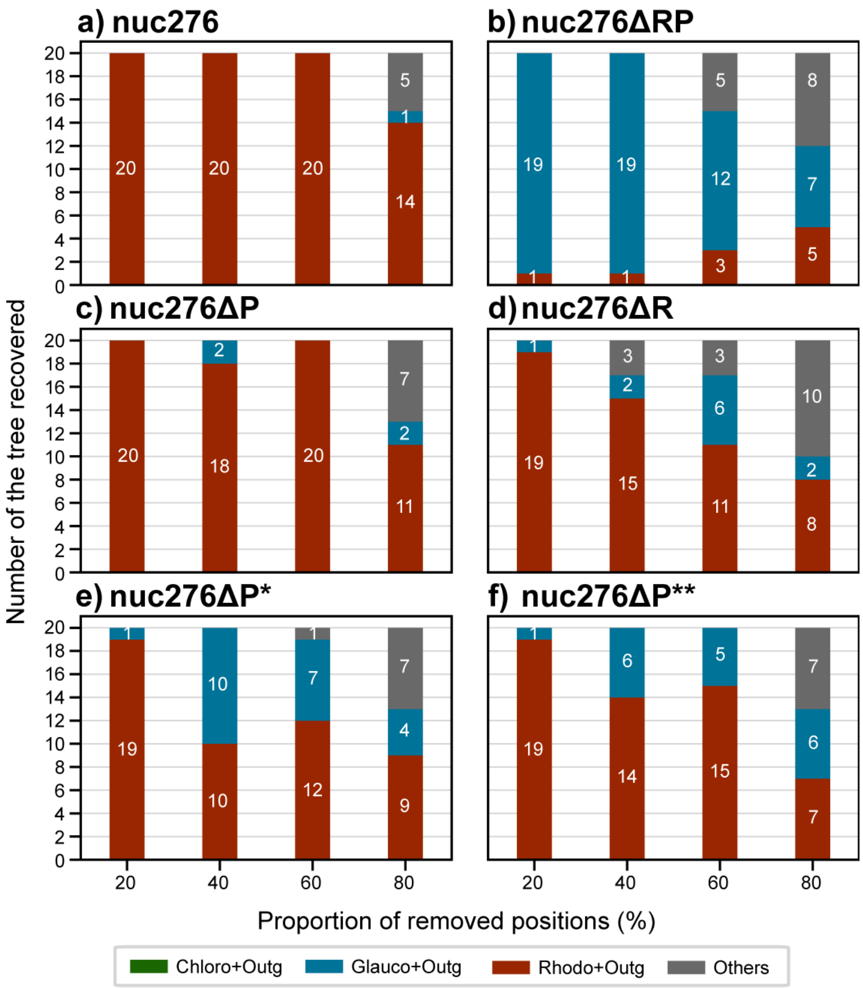


**Supplementary figure 6. The frequency of tree topologies inferred from nuc276 (a), nuc276ΔRP (b), nuc276ΔP (c), nuc276ΔR (d), nuc276ΔP* (e), and nuc276ΔP** (f) processed by random position removal (RPR) analysis.** The frequencies of the competing tree topologies for the internal relationship among Chloroplastida, Glaucophayta, and Rhodozoa recovered from the RPR-processed supermatrices. For each data point, we obtained 20 trees inferred from the RPR-processed supermatrices and counted the occurrences of the three trees, which are different in the basal sub-clade in the Archaeplastida clade. In the bar graphs, green represents the occurrence of the "Chloro+Outg" tree, in which the sister relationship between Chloroplastida and the outgroup, was recovered (see the inset). Likewise, the occurrences of the "Glauco+Outg" and "Rhodo+Outg" trees, in which Glaucophyta and Rhodozoa were tied with the outgroup, are represented by blue and red, respectively. The three types of the tree of Archaeplastida satisfied the criteria in which Chloroplastida, Glaucophyta, Rhodozoa, Pancryptista, and the six other eukaryotes formed individual clades, in addition to the former three grouped together (i.e., the monophyly of Archaeplastida). Trees, which dissatisfied with any of the above criteria, were designated as "Others."

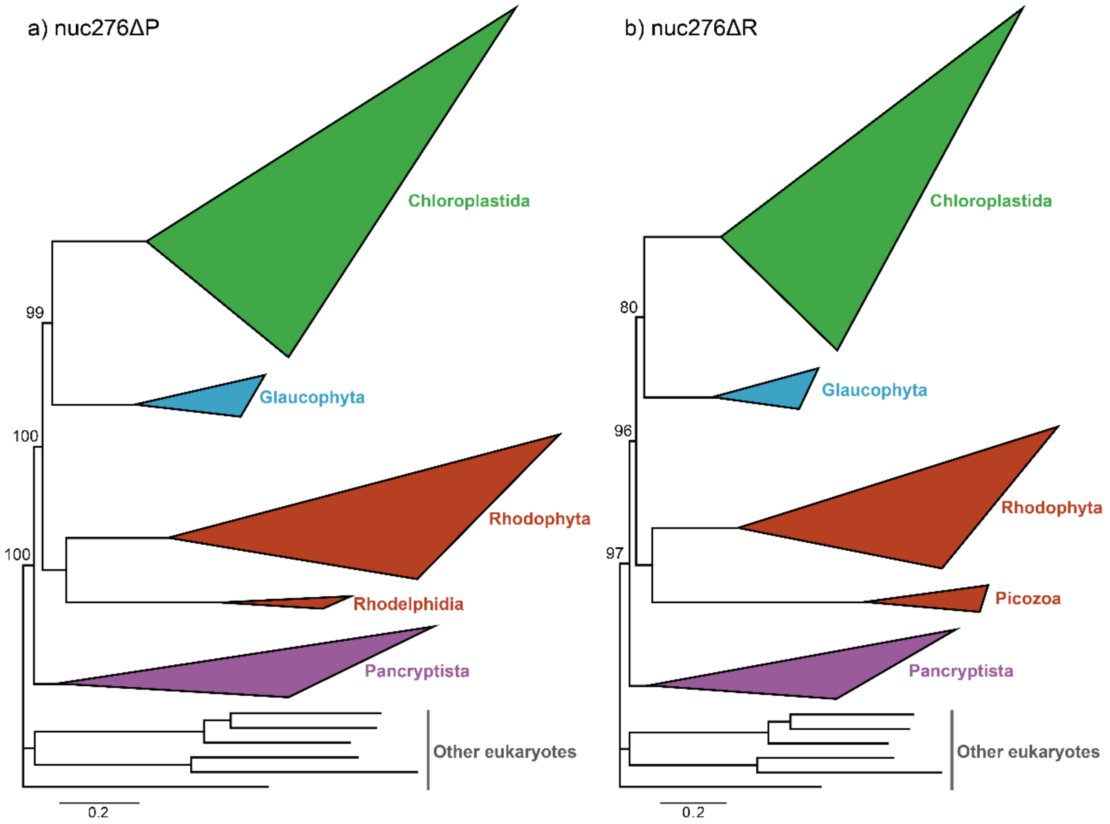


**Supplementary figure 7. The trees of Archaeplastida inferred from nuc276ΔP and nuc276ΔR.** The phylogenetic trees were recovered by the maximum likelihood method. The trees inferred from nuc276ΔP and that from nuc276ΔR are shown in (a) and (b), respectively. The three sub-clades in Archaeplastida were simplified into triangles. The size of each triangle is scaled to the number of the OTUs comprising the clade, and the longest and shortest distances from the ancestral to terminal nodes. All OTU names were omitted. We also omitted the support values for all bipartitions, except the ultrafast bootstrap percent values for the monophyly of Archaeplastida and the basal branching of Rhodophyta plus Rhodelphidia or Picozoa.

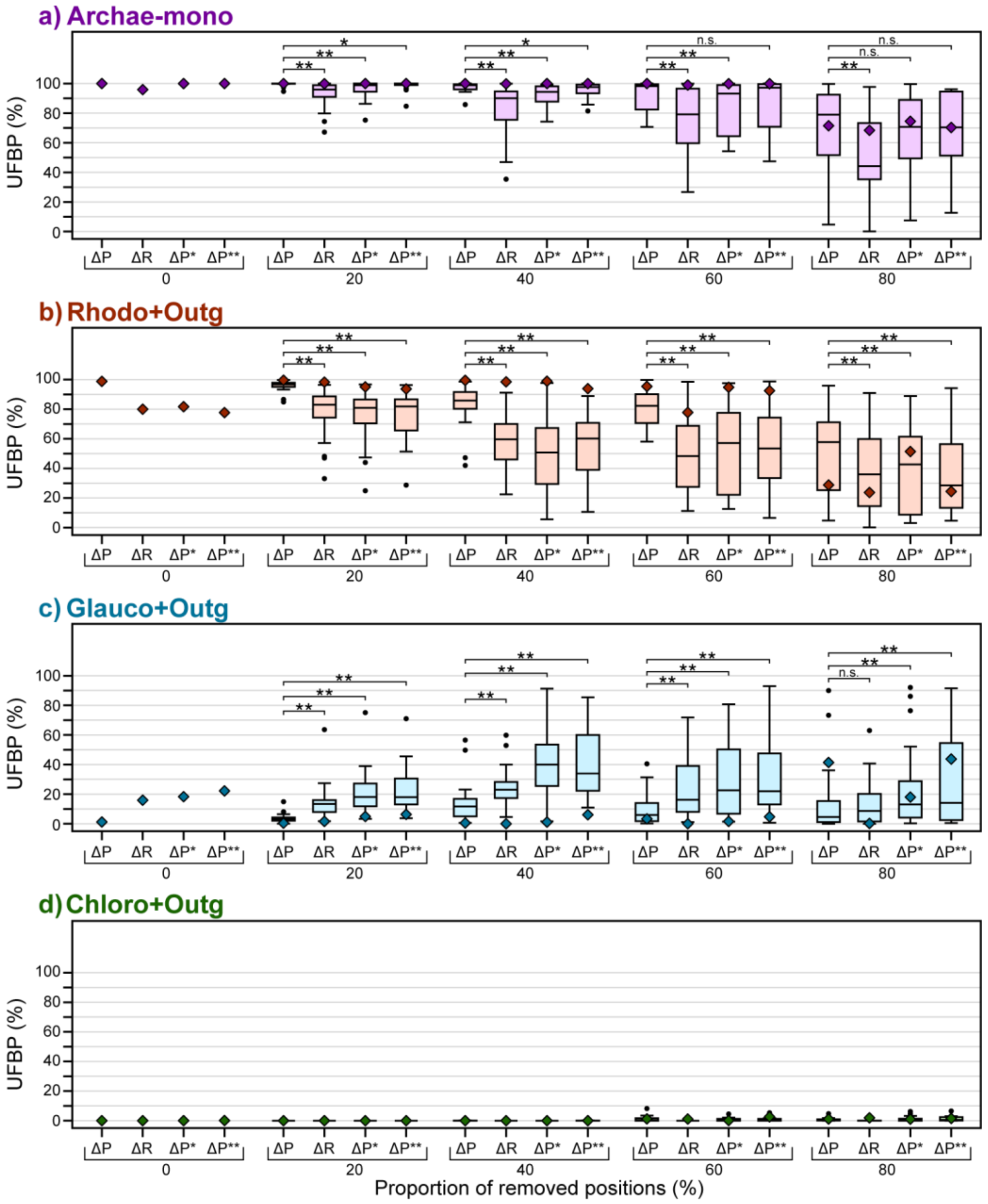


**Supplementary figure 8. Analyses of nuc276ΔP, nuc276ΔR, nuc276ΔP*, and nuc276ΔP** processed by fast-evolving position removal (FPR) and random position removal (RPR) analyses.** (a) UFBPs for the monophyly of Archaeplastida. (b) UFBPs for the Rhodo+Outg tree in which Rhodozoa is excluded from the union of Chloroplastida and Glaucophyta in the Archaeplastida clade. (c) UFBPs for the Glauco+Outg tree in which Glaucophyta is excluded from the union of Chloroplastida and Rhodozoa in the Archaeplastida clade. (d) UFBPs for the Chloro+Outg tree in which Chloroplastida is excluded from the union of Rhodozoa and Glaucophyta in the Archaeplastida clade. The details of this figure are the same as those of Figure 3, except that ultrafast bootstrap percent values (UFBPs) calculated from the FPR-processed supermatrices are shown in diamonds. The UFBPs calculated from the RPR-processed nuc276ΔP and nuc276ΔR (or nuc276ΔP* or nuc276ΔP**) were compared with the Wilcoxon signed-rank test. Double and single asterisks indicate that the null hypothesis of no difference in the median values of UFBPs from the two analyses was rejected with $p$ <0.05 and <0.05, respectively. n.s.; not significant ($p$ ≧0.05).

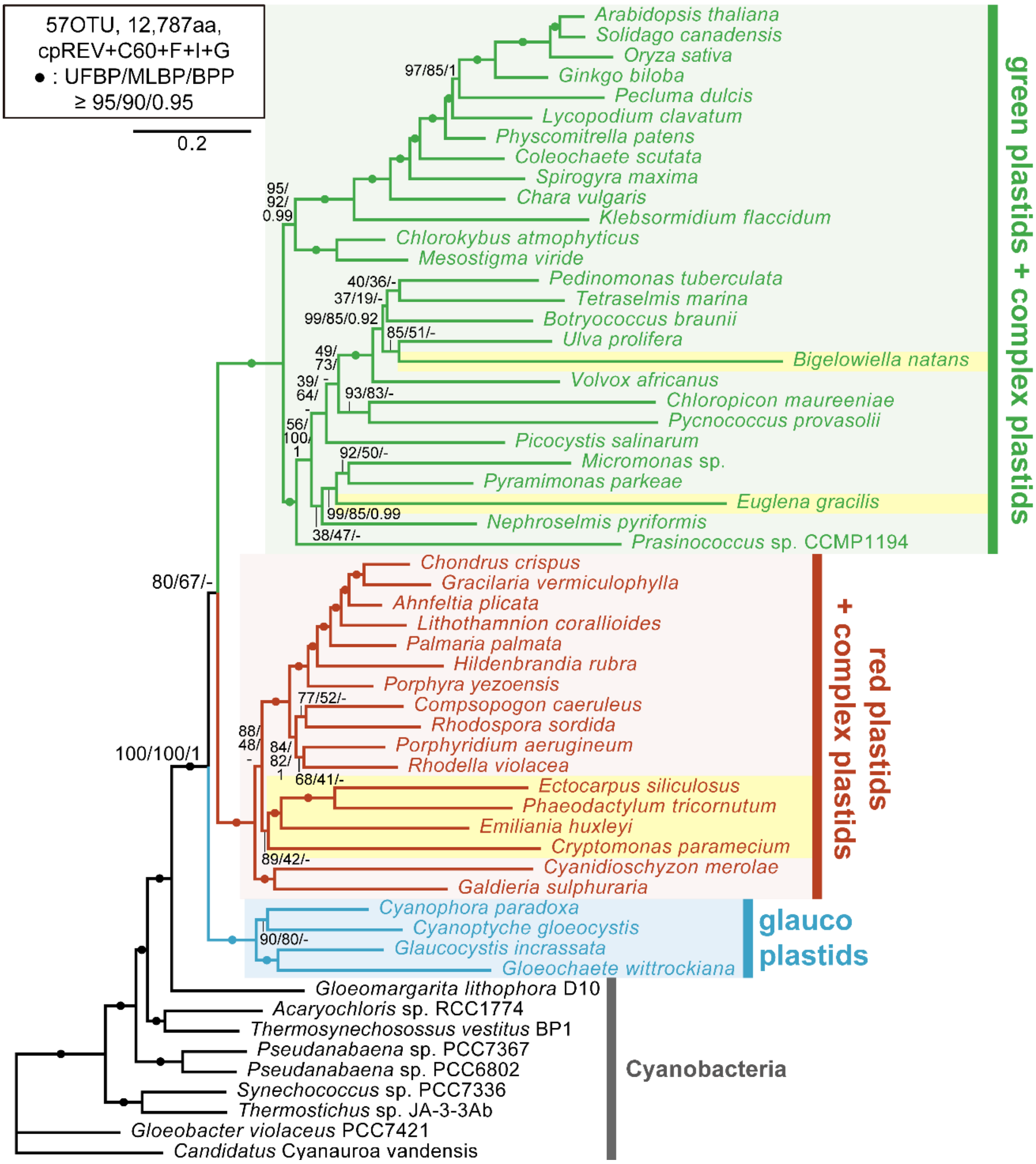


**Supplementary figure 9. The ML tree of plastids inferred from pld54.** The phylogenetic tree was recovered by the maximum likelihood (ML) method. For each bipartition, an ultrafast bootstrap percent value (UFBP), ML bootstrap values (MLBPs), and a Bayesian posterior probability (BPP) are shown on the left, center, and right, respectively. The bipartitions marked by dots received UFBPs ≧95%, MLBPs ≧90%, and BPPs ≧0.95. If the ML and Bayesian analyses recovered identical bipartitions, BPPs were shown along with the corresponding UFBPs and MLBPs in the figure. For the details of the Bayesian consensus tree, see Fig. S11. The OTUs corresponding to the six complex plastids are shaded in yellow.

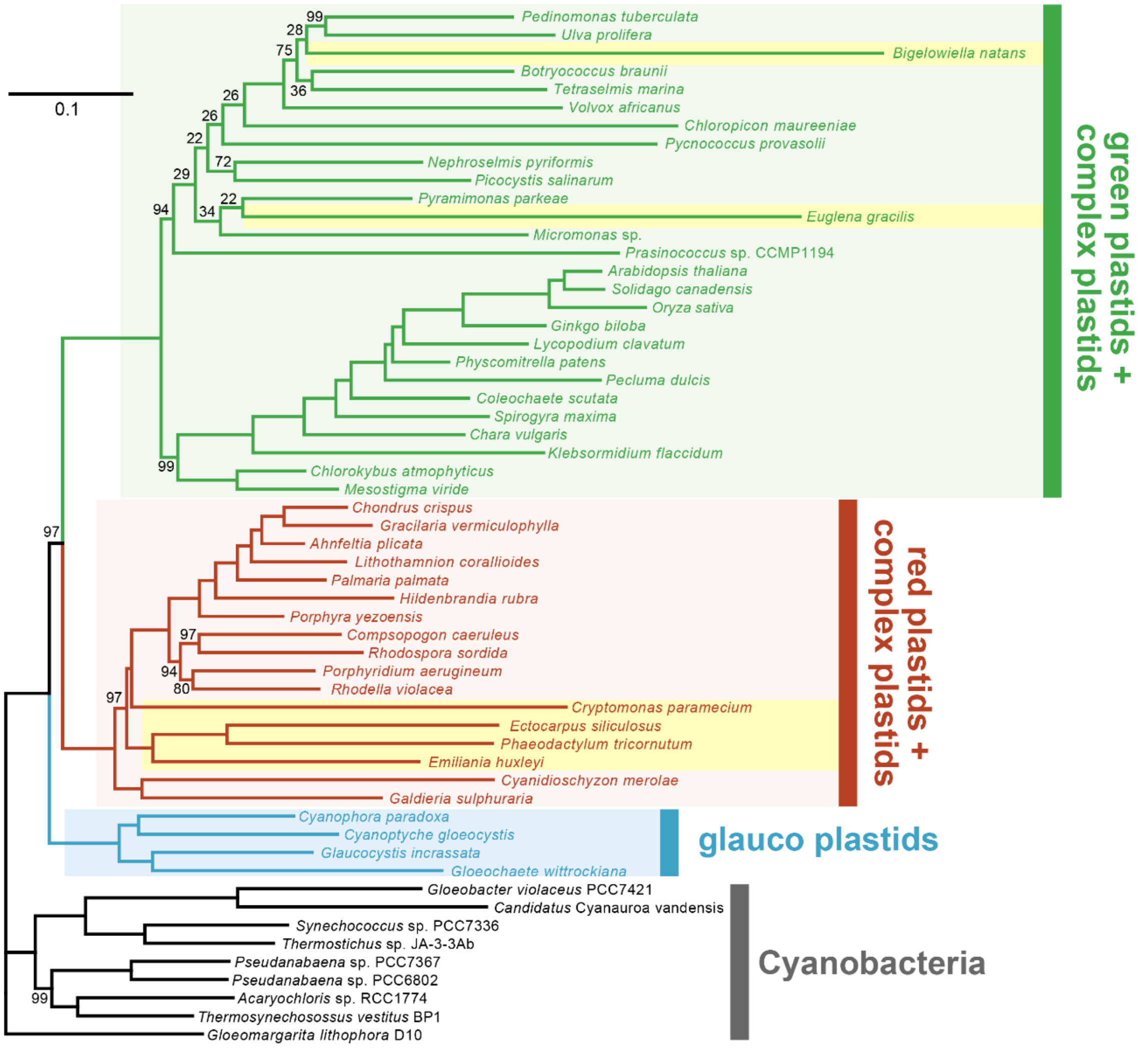


**Supplementary figure 10. The ML tree of plastids inferred from pld54 after 3,152 positions bearing potential bias in amino acid composition**. The details of this figure are the same as those of Figure S9. For all bipartitions, UFBPs are presented except for those with 100%.

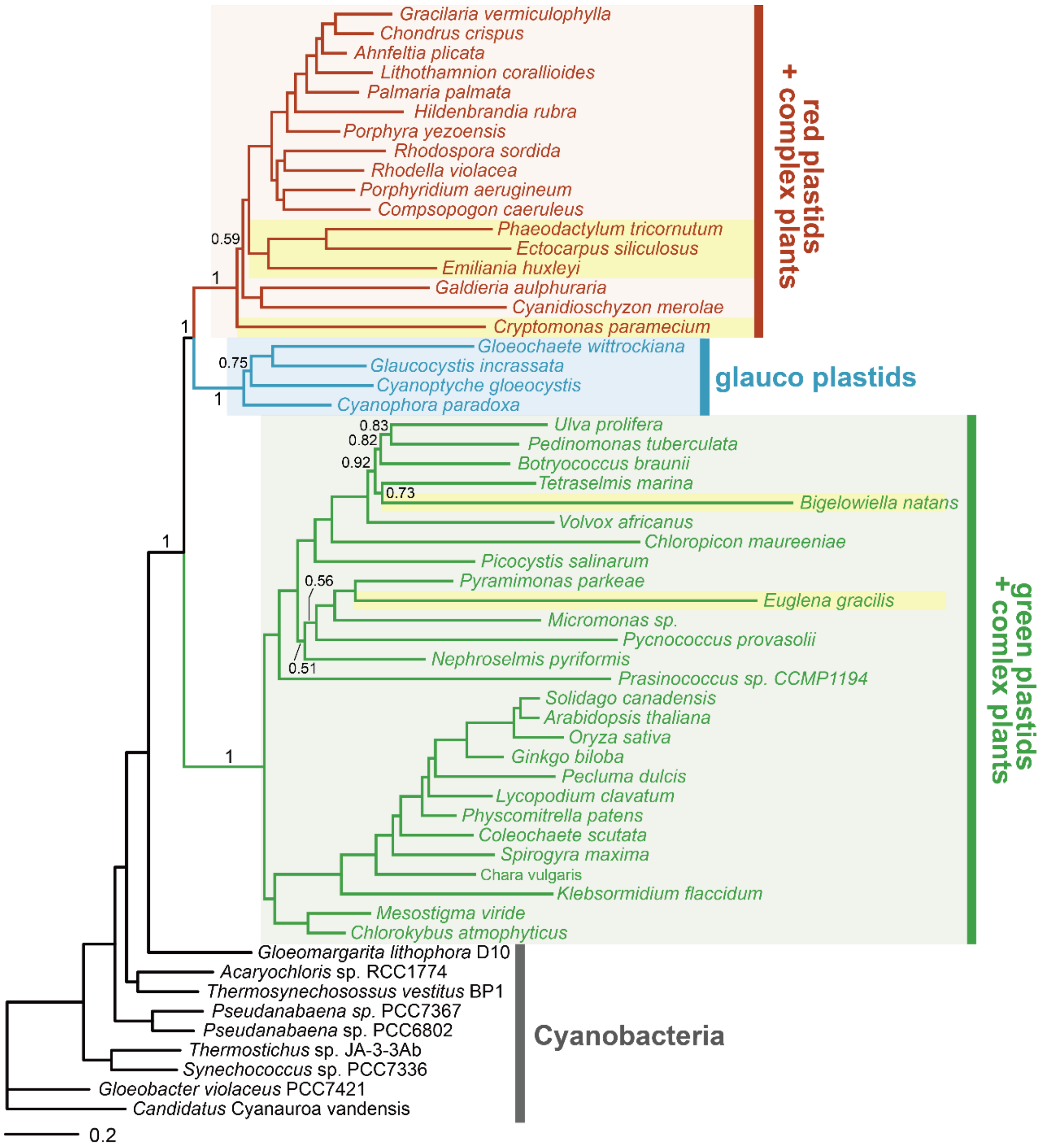


**Supplementary figure 11. Bayesian tree of plastids inferred from pld54**. The details of this figure are the same as those of Figure S9. The BPPs ≧0.95 are omitted, except for those for the nodes important for the relationship among the three types of primary plastids.

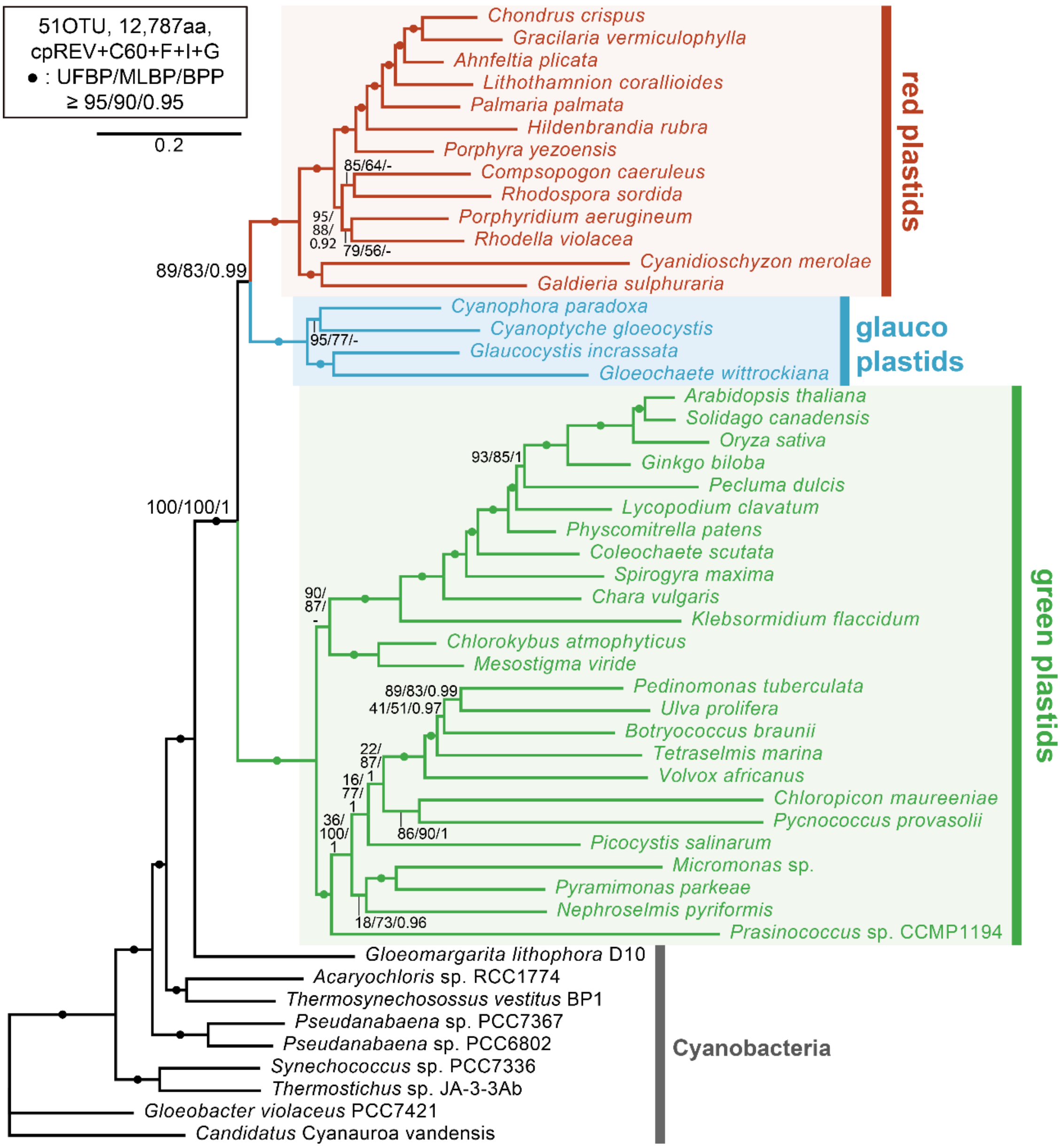


**Supplementary figure 12. The ML tree of plastids inferred from pld54Δg*r*.** The details of this figure are the same as those of Supplementary Figure 9. Note that the relationship among green plastids, glaucophyte plastids, and red plastids inferred from the ML and Bayesian analyses agreed with each other.

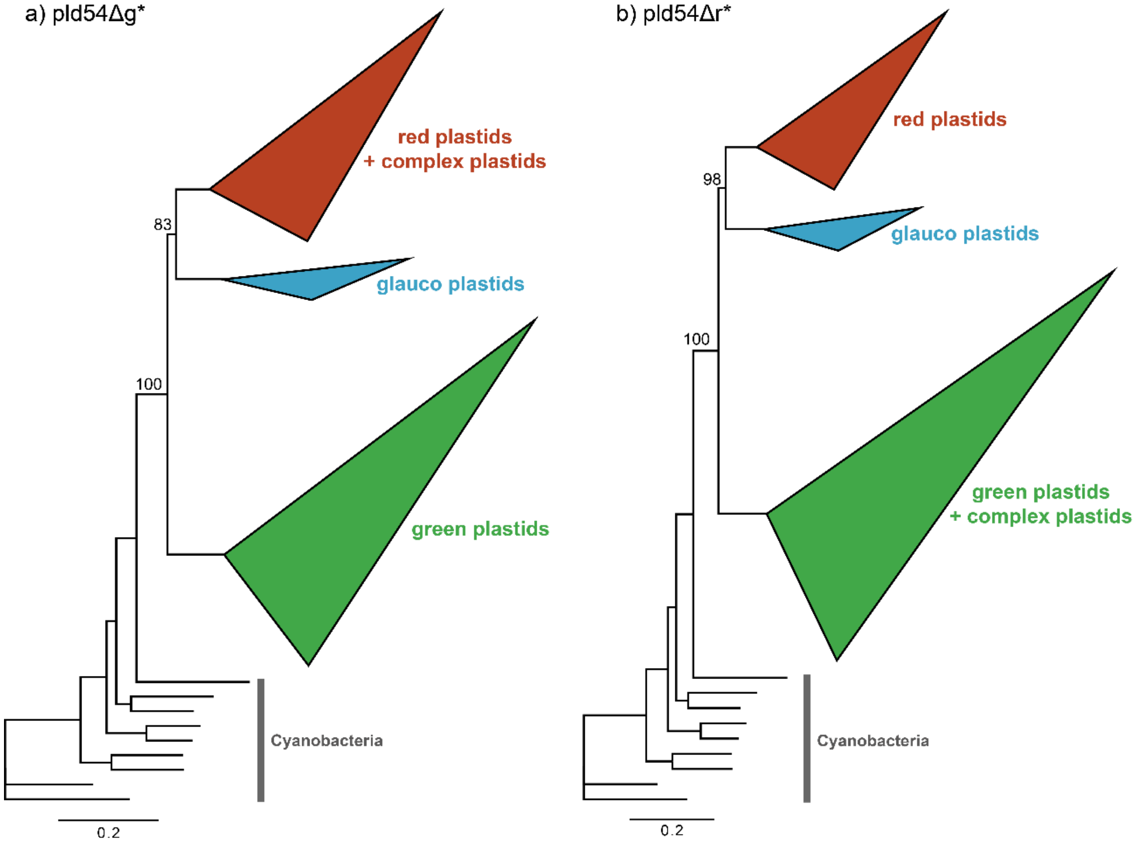


**Supplementary figure 13. The trees of plastids inferred from pld54Δg* and pld54Δr*.** The phylogenetic trees were recovered by the maximum likelihood (ML) method. The trees inferred from pld54Δg* and that from pld54Δr* are shown in (a) and (b), respectively. The three sub-clades in the plastid clade were simplified into triangles. The size of each triangle is scaled to the number of the OTUs comprising the clade, and the longest and shortest distances from the ancestral to terminal nodes. All OTU names were omitted. We also omitted the support values for all bipartitions, except the ultrafast bootstrap percent values for the monophyly of plastid clade and the basal branching of green plastids (and green complex plastids in the tree from pld54Δr* in b).

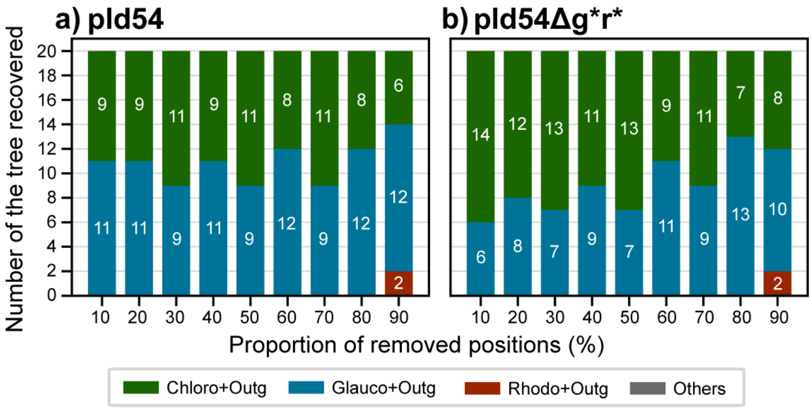


**Supplementary figure 14. The frequency of tree topologies inferred from pld54, pld54Δg*r* processed by random position removal (RPR) analysis.** The frequencies of the competing tree topologies for the internal relationship among green plastids, glaucophyte plastids, and red plastids recovered from the RPR-processed pld54 are shown in (a). The results from the same analyses based on pld54Δg*r* are shown in (b). The plot details of this figure are the same as those of the supplementary figure 6. About the criteria of topologies, the three types of the tree of plastids satisfied the criteria in which green plastids with or without green alga-derived complex plastids, glauco plastids, red plastids with or without red alga-derived complex plastids, and Cyanobacteria formed individual clades, in addition to the former three grouped together (i.e., the monophyly of plastids). Trees, which dissatisfied with any of the above criteria, were designated as “Others.”

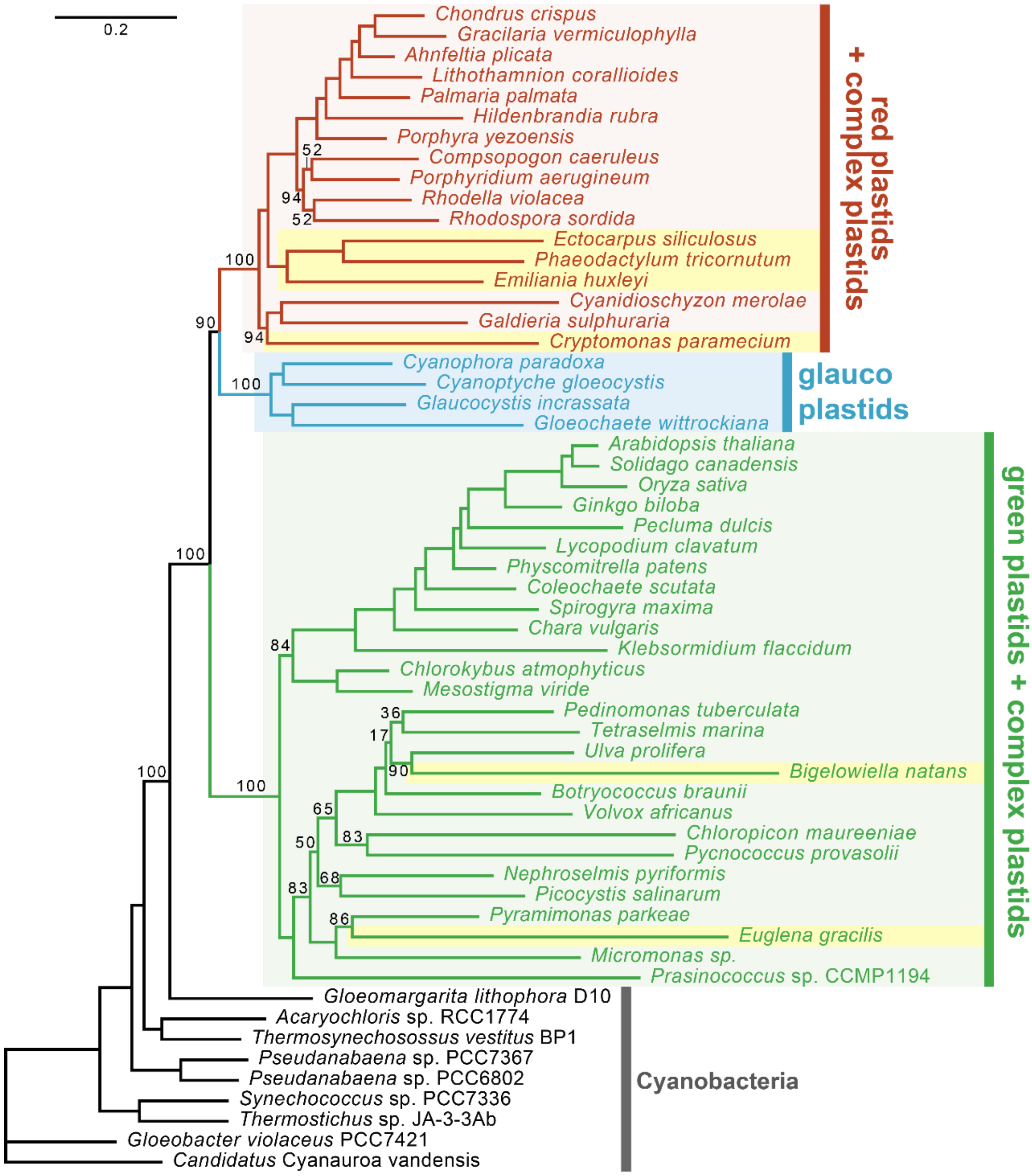


**Supplementary figure 15. The ML tree of plastids inferred from pld54 after removing the alignment positions with the 5% largest and 5% smallest site-rate ratios**. The details of this figure are the same as those of Figure S9. The UFBPs ≧95% were omitted, except for those for the nodes important for the relationship among the three types of primary plastids.
